# *Drosophila* tachykininergic neurons modulate the activity of two groups of receptor-expressing neurons to regulate aggressive tone

**DOI:** 10.1101/2021.10.11.463893

**Authors:** Margot Wohl, Kenta Asahina

## Abstract

Neuropeptides influence animal behaviors through complex molecular and cellular mechanisms, many of which are difficult to predict solely from synaptic connectivity. Here, we uncovered two separate downstream targets that are differentially modulated by the neuropeptide tachykinin, which promotes *Drosophila* aggression. Tachykinin from a single sexually dimorphic group of neurons recruits two separate downstream groups of neurons. One downstream group, synaptically connected to the tachykinergic neurons, expresses the receptor *TkR86C* and is necessary for aggression. Tachykinin supports the strength of cholinergic excitatory synaptic transmission between the tachykinergic and *TkR86C* downstream neurons. The other downstream group expresses the *TkR99D* receptor and is recruited primarily when tachykinin is over-expressed in the source neurons. This circuit reconfiguration correlates with the quantitative and qualitative enhancement of aggression observed when tachykinin is present in excess. Our data highlight how the amount of neuropeptide released from a small number of neurons can reshape the activity patterns of multiple downstream neural populations.

## INTRODUCTION

Neuromodulation plays an important role in controlling ethologically important “survival behaviors” (Castro and Bruchas, 2019; LeDoux, 2012), including social behaviors (Insel, 2010). Neuropeptides are a major class of neuromodulator important for a variety of innate behaviors, such as feeding, fear and stress responses, sleep, and reproduction (Castro and Bruchas, 2019; Nässel and Winther, 2010). Because of its behavioral relevance, the neuropeptidergic system has been a major target for developing effective therapeutics (Griebel and Holsboer, 2012; Hökfelt et al., 2003; Holmes et al., 2003). While neuropeptides that are released into the circulatory system act as neurohormones, growing evidence indicates that neuropeptides can also locally modulate specific target neurons (Nässel, 2009; Nusbaum et al., 2017; Salio et al., 2006; van den Pol, 2012). For instance, several neuropeptides are known to alter the physiology of a critical circuit node only during a specific hunger state, which ultimately changes the dynamics of the behavior-controlling circuit (Ko et al., 2015; Krashes et al., 2009; Oh et al., 2019). Flexibility in release sites and co-transmission with fast-acting neurotransmitters means that neuropeptides can impact the physiology of neurons beyond that predicted by the connectome (Bargmann, 2012; Marder, 2012; Nässel, 2009; Nusbaum et al., 2017; Salio et al., 2006; van den Pol, 2012). Indeed, findings in invertebrate nervous systems, such as those of crustaceans and nematodes, indicate that the behaviorally relevant “chemoconnectomes” of neuromodulators are dynamic and multifunctional (Flavell et al., 2013; Leinwand and Chalasani, 2013; Nusbaum et al., 2017). Although specific neuropeptidergic cell populations are often important for controlling survival behaviors in both vertebrates and invertebrates, how a single source of neuropeptides can coordinate the activity of multiple behaviorally relevant target neurons remains poorly understood.

In this study, we characterized the impacts of peptidergic neuromodulation in microcircuits that control inter-male aggression in the fruit fly, *Drosophila melanogaster*. The male-specific Tk-GAL4^FruM^ neurons are known to promote aggressive behavior in part by releasing neuropeptide tachykinin (Asahina et al., 2014; Wohl et al., 2020). We created new genetic alleles that label tachykinin receptor-expressing neurons to probe how tachykinin modulates targets downstream of Tk-GAL4^FruM^ neurons. Functional calcium imaging across the brain revealed two distinct, spatially restricted subsets of downstream neurons, each expressing a different *Drosophila* tachykinin receptor (*TkR86C* or *TkR99D*). Tachykininergic modulation of these targets by Tk-GAL4^FruM^ activation differed both quantitatively and qualitatively and was correlated with the level and specificity of aggressive behaviors. Collectively, our results identify a receptor-based neuronal mechanism of tachykininergic neuromodulation. This example provides insight into how neuropeptides can act to control a complex behavior by reshaping the physiological dynamics of a target circuit, underscoring the significance of functional connectivity based on peptide–receptor relationships (the “chemoconnectome”).

## RESULTS

### Tachykinins in Tk-GAL4^FruM^ neurons quantitatively and qualitatively enhance aggression

Tk-GAL4^FruM^ neurons promote aggression toward other males, but not toward females, likely due to a *doublesex* (*dsx*)-dependent mechanism that enforces target specificity of male aggression (Wohl et al., 2020). Previous work that used a thermogenetic approach did not address whether tachykinin released from Tk-GAL4^FruM^ neurons can alter the target sex specificity of male aggression (Asahina et al., 2014). Here, we quantified male- and female-directed aggressive behavior induced by optogenetic activation of Tk-GAL4^FruM^ neurons by the red-shifted channelrhodopsin CsChrimson (Klapoetke et al., 2014), while varying the level of tachykinin expression in these neurons (Fig. 1A).

**Figure 1:**
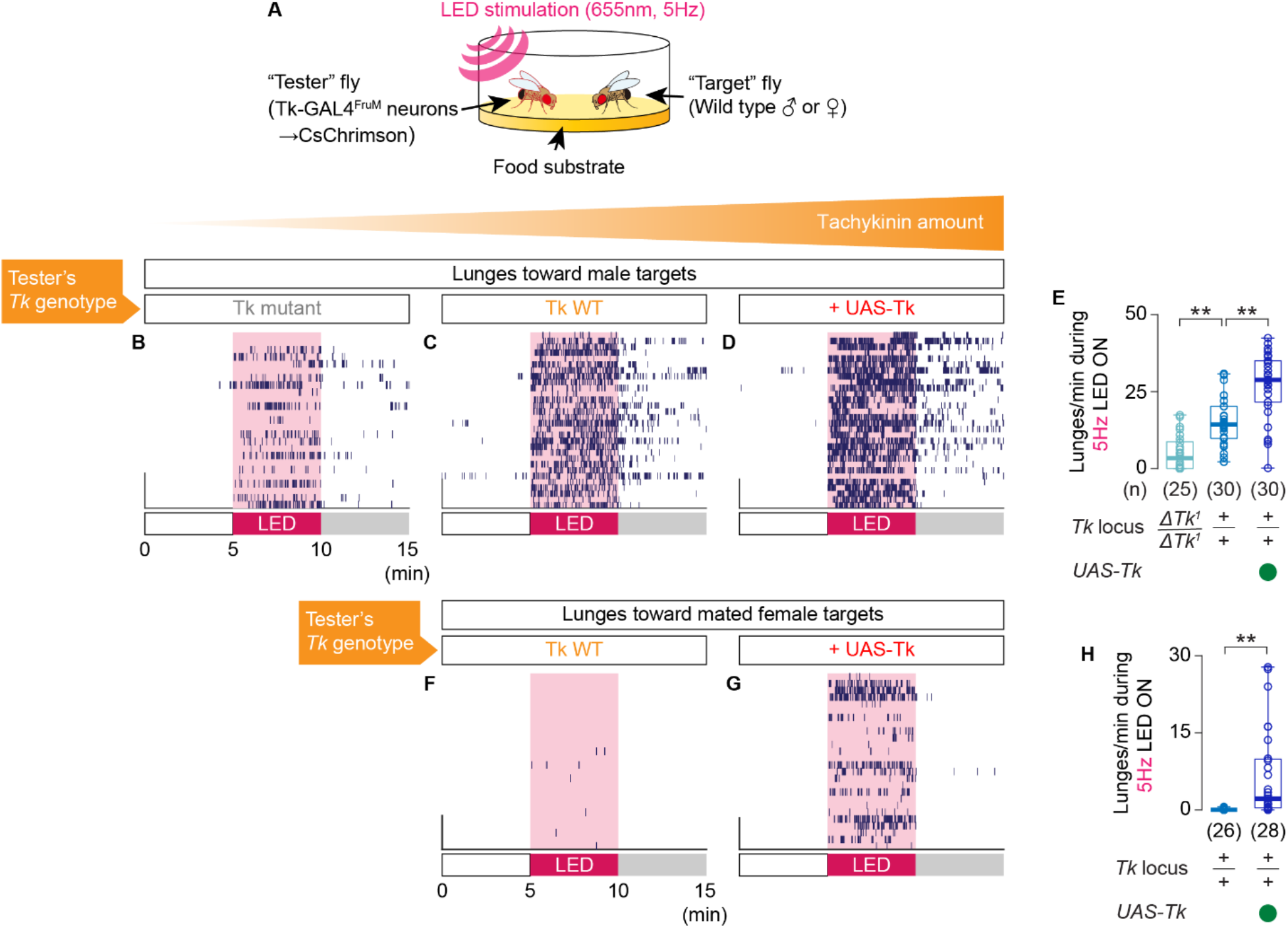
Tachykinin amount controls the intensity of aggression induced by optogenetic activation of Tk-GAL4^FruM^ neurons. **A.** Design of the optogenetic behavioral assay. **B-D, F-G.** Raster plots of lunges toward a male (**B-D**) or a mated female (**F**, **G**) target fly induced by optogenetic activation of Tk-GAL4^FruM^ neurons. **E**, **H**. Boxplots of male- (**E**) and female- (**H**) targeted lunges performed during optogenetic stimulation of Tk-GAL4^FruM^ neurons. In **E**, ** p < 0.01 by Kruskal-Wallis one-way ANOVA (p < 2.3X10^−10^) and post-hoc Mann-Whitney U-test. In **H**, ** p < 0.01 by Mann-Whitney U-test. **B** is data from *Tk* null mutants, **C** and **F** are data from *Tk* wild type, and **D** and **G** are data from animals with a *UAS-Tk* transgene. In **E** and **H**, the *Tk* genotypes are indicated at the bottom.

Consistent with the results from thermogenetic manipulation, the tachykinin null mutation attenuated male-directed aggression induced by optogenetic activation of Tk-GAL4^FruM^ neurons, whereas over-expression of tachykinin in Tk-GAL4^FruM^ neurons enhanced male-directed aggression, at two different stimulation frequencies (Fig. 1B-E, Fig. S1A-E). Heterozygosity of the *Tk* null mutation did not affect the frequency of lunges induced by optogenetic activation of Tk-GAL4^FruM^ neurons (Fig. S1D, E), consistent with the previous observation that the expression level of *Tk* remained unchanged in the *Tk* null heterozygotes (Asahina et al., 2014). Aggression levels were comparably low among genotypes during the pre-stimulation time windows (Fig. S1F, G), suggesting that tachykinin needs to be released in an activity-dependent manner to promote aggression. Also, over-expression of tachykinin did not increase persistent aggression in the post-stimulus time window (Fig. S1F, H), further arguing that tachykinin promotes aggression by enhancing the immediate physiological impact of Tk-GAL4^FruM^ neuronal activity on the circuit.

Intriguingly, over-expression of tachykinin caused male tester flies to attack female targets during optogenetic stimulation, which was rare in wild-type flies (Fig. 1F-H). Such qualitative enhancement of aggression may be mediated by recruitment of a new circuit component. These results suggest that tachykinin from Tk-GAL4^FruM^ neurons is involved in both quantitative (toward males) and qualitative (toward females) enhancement of male aggressive behavior.

### Anatomical relationship between *TkR86C*-expressing neurons and Tk-GAL4^FruM^ neurons

To begin elucidating the downstream targets of Tk-GAL4^FruM^ neurons, we first needed to identify which arborizations were dendritic and which were axonal. We used the genetically encoded post-synaptic marker DenMark (Nicolaï et al., 2010) to identify dendrites, and the pre-synaptic marker synaptotagmin:GFP (Syt:GFP) (Zhang et al., 2002) to identify axon terminals of Tk-GAL4^FruM^ neurons. Post-synaptic (dendritic) markers were primarily detected in arborizations in the lateral crescent, ring, and lateral junction structures (Cachero et al., 2010; Yu et al., 2010) (Fig. 2A), which are proposed to integrate olfactory and gustatory information (Auer and Benton, 2016; Clowney et al., 2015; Yu et al., 2010). On the other hand, pre-synaptic markers were primarily detected in the branches projecting to the superior medial protocerebrum (SMP) and in the bilateral arch (Ito et al., 2014; Yu et al., 2010) (Fig. 2B), which were largely devoid of DenMark signal (Fig. 2A_2,3_). Both pre- and post-synaptic markers were mostly undetectable in the commissural tract that extends from the dorsal side of the lateral junction (Fig. 2A_2,3_, B_2,3_). The Syt:GFP-enriched branches to the SMP emanate from this tract, suggesting that it is the axonal tract of Tk-GAL4^FruM^ neurons.

**Figure 2:**
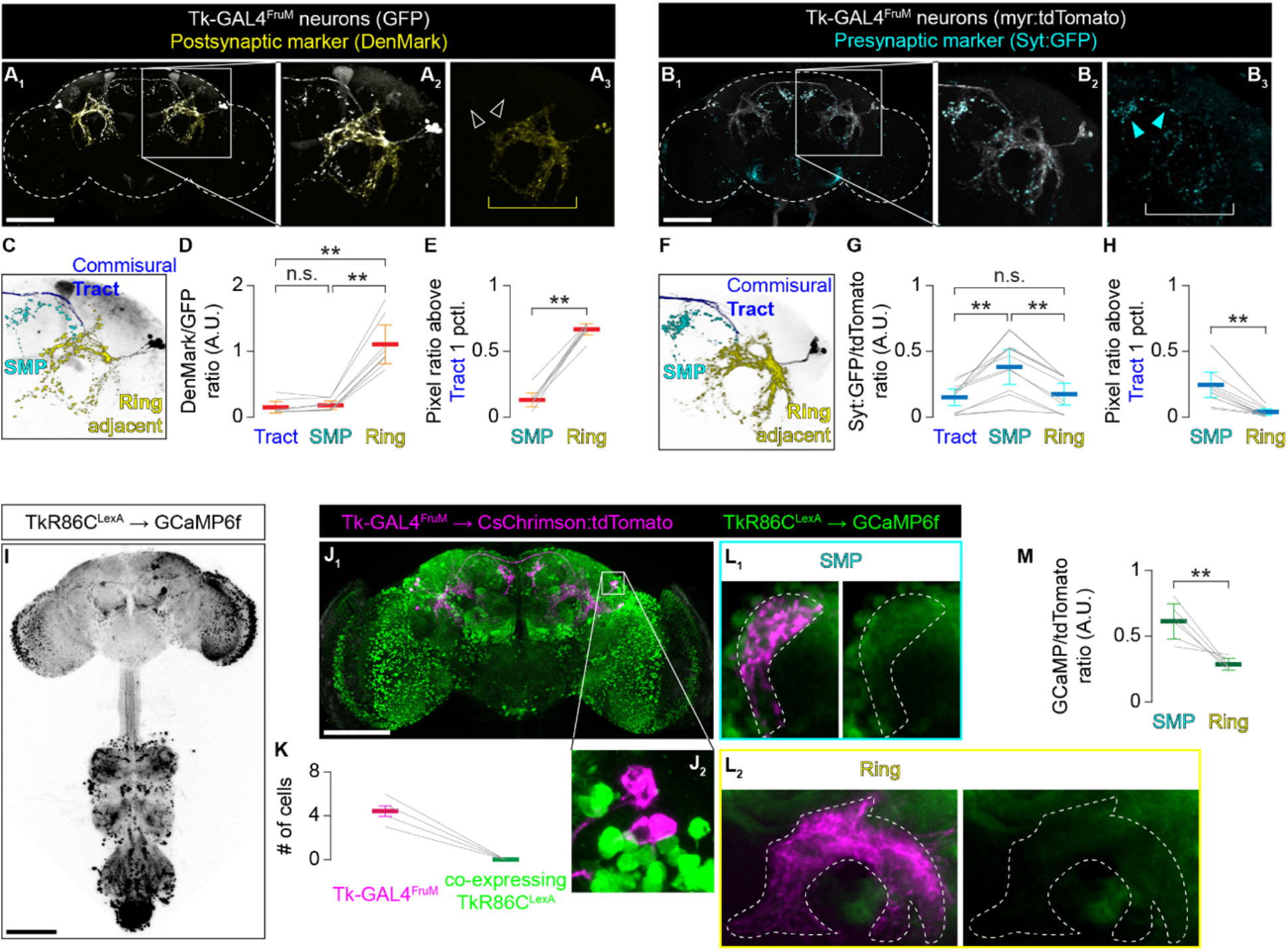
Tk-GAL4^FruM^ neurons are positioned to make synaptic contact with *TkR86C^LexA^* neurons. **A.** A representative image of a male brain expressing GFP (white) and the postsynaptic marker DenMark (yellow), which is present in the ring-adjacent region (yellow bracket in **A3**) but not in the projection to the SMP (empty white arrowheads). Scale bar: 100 µm. **B.** A representative image of a male brain expressing myristoylated tdTomato (myr:tdTomato, white) and the presynaptic marker synaptotagmin:GFP (Syt:GFP, cyan), which is present in the projection to the SMP (cyan arrowheads in **B3**) but only sparsely observed in the ring-adjacent region (white bracket). Scale bar: 100 µm. **C, F.** Segmentation of the region shown in **A2** (**C**) and **B2** (**F**). **D, G**. DenMark (**D**, n = 8) and Syt:GFP (**G**, n = 12) immunohistochemical signals relative to GFP (**D**) and myr:tdTomato (**G**) signals in the commissural tract (tract), SMP, and ring-adjacent region (ring). ** p < 0.01, n.s. p > 0.05 by two-way ANOVA (**D**: p 2.0 × 10^−8^, **G**: p < 1.2 × 10^−6^) and post-hoc Tukey’s Honest Significance test. **E, H.** DenMark (**E**, n = 8) and Syt:GFP (**H**, n = 12) immunohistochemical signals in SMP and ring areas relative to the signals in the tract. ** p < 0.01 by paired t-test. **I.** Representative expression pattern of GCaMP6f driven by *TkR86C^LexA^* in the nervous system, visualized by immunohistochemistry. Scale bar: 100 µm. **J.** A representative image of a male brain expressing GCaMP6f driven by *TkR86C^LexA^* (green) and CsChrimson:tdTomato under intersectional control of *Tk-GAL4^1^* and *fru^FLP^* (magenta). Scale bar: 100 µm. **K.** *TkR86C^LexA^* does not label Tk-GAL4^FruM^ neurons (n = 12). **L.** Distribution of immunohistochemical signals of GCaMP6f driven by *TkR86C^LexA^* (green), and CsChrimson:tdTomato under intersectional control of *Tk-GAL4^1^* and *fru^FLP^* (magenta). The magnified images near the Tk-GAL4^FruM^ neuronal projections (broken white lines) in the SMP (**L1**) and the ring region (**L2**) from an averaged image stack of eight standardized hemibrains (see Materials and Methods for details) are shown. **M.** Average GCaMP6f immunohistochemical fluorescence in the SMP and ring-adjacent region as defined by CsChrimson:tdTomato immunohistochemical signals in Tk-GAL4^FruM^ neurons. ** p < 0.01 by paired t-test. The thick line and error bars in **D**, **E**, **G**, **K, H**, and **M** represent the average and 95% confidence intervals.

To quantify these observations, we segmented Tk-GAL4^FruM^ neurons into three domains: arborizations in the SMP, arborizations in lateral regions (hereafter called “ring adjacent” regions), and the commissural tracts (Fig. 2C, F). We then measured the average signal intensity of both Syt:GFP and DenMark within each domain. As expected, DenMark signals were enriched in the ring adjacent region (Fig. 2D, E) while Syt:GFP signals were enriched in the SMP projection (Fig. 2G, H). Punctated Syt:GFP signals were also sparsely detected in regions of the Tk-GAL4^FruM^ neurons enriched with DenMark signals (Fig. 2B_2,3_). At least some of this Syt:GFP signal likely belongs to pre-synaptic termini from the contralateral projection. Samples from brains with Tk-GAL4^FruM^ neurons labeled unilaterally show that the axonal commissural tract crosses the midline and projects to a medial part of the ring on the contralateral side (Fig. S2A, B). It is also possible that the ring adjacent region contains presynaptic sites that mediate retrograde or dendro-dendritic communications. Overall, these largely segregated distributions of pre- and post-synaptic markers suggest that neurotransmitters from Tk-GAL4^FruM^ neurons are mainly released in the SMP.

Previous work showed that mutation of the tachykinin receptor gene *TkR86C* attenuates aggression triggered by thermogenetic excitation of Tk-GAL4^FruM^ neurons (Asahina et al., 2014). This suggests that at least a subset of the circuit downstream of Tk-GAL4^FruM^ neurons expresses *TkR86C*. To visualize these putative downstream neurons, we created a novel knock-in allele of *TkR86C*, named *TkR86C^LexA^*, using CRISPR/Cas9-mediated gene editing (Gratz et al., 2014) (Fig. S2C). *TkR86C^LexA^*-expressing neurons were numerous and widespread (visualized with immunohistochemistry against LexA-driven GCaMP6f (Chen et al., 2013)), both in the central brain and in the ventral nerve cord (Fig. 2I). This expression pattern is similar to that of a previously reported *TkR86C* knock-in allele (Kondo et al., 2020). The *TkR86C^LexA^* expression pattern is also consistent with the broad expression of tachykinin peptides (Winther et al., 2003). Importantly, Tk-GAL4^FruM^ neurons do not express *TkR86C^LexA^* (Fig. 2J_1-2_, K), suggesting that tachykininergic modulation by Tk-GAL4^FruM^ neurons through *TkR86C* does not employ an autocrine mechanism (Choi et al., 2012).

We next asked whether *TkR86C*-expressing neurons and Tk-GAL4^FruM^ neurons are directly connected by examining the anatomical relationship between these two neuronal populations. Immunohistochemistry revealed that the pre-synaptic regions of Tk-GAL4^FruM^ neurons in the SMP are in close proximity to the neuronal processes of *TkR86C^LexA^* neurons (Fig. 2L_1_). By contrast, *TkR86C^LexA^* neurons showed less overlap with the post-synaptic ring-adjacent regions of Tk-GAL4^FruM^ neurons (Fig. 2L_2_, M). This suggests that some *TkR86C^LexA^* neurons are positioned to receive synaptic inputs in the SMP from Tk-GAL4^FruM^ neurons.

### *TkR86C*-expressing neurons are functionally downstream of Tk-GAL4^FruM^ neurons

We next sought to obtain physiological evidence that *TkR86C^LexA^* neurons receive neural input from Tk-GAL4^FruM^ neurons. The anatomical results thus far are consistent with the idea that a subset of *TkR86C^LexA^* neurons is synaptically downstream of Tk-GAL4^FruM^ neurons. However, the mere proximity of neurites does not guarantee the presence of synapses. Moreover, while some studies have observed peptide-containing dense core vesicles primarily near presynaptic sites (Jan et al., 1980; Salio et al., 2006; Schlegel et al., 2016; Tao et al., 2018), neuropeptides are also released extrasynaptically (Jan and Jan, 1982; Karhunen et al., 2001) and affect the physiology of target neurons that are not synaptically connected (Jan and Jan, 1982; Jan et al., 1980; Nässel, 2009; van den Pol, 2012). To determine whether *TkR86C^LexA^* neurons receive neural input from Tk-GAL4^FruM^ neurons near their synaptic termini or in extra-synaptic locations, we visualized *TkR86C^LexA^* neuronal activity patterns across a large portion of the brain in response to optogenetic excitation of Tk-GAL4^FruM^ neurons.

We created a fly that expressed CsChrimson specifically in Tk-GAL4^FruM^ neurons and the genetically encoded calcium indicator GCaMP6f specifically in *TkR86C^LexA^* neurons. We used two-photon serial volumetric imaging to monitor the fluorescence intensity of GCaMP6f in multiple z-planes (dorsal to ventral) of the brain in live flies (Siju et al., 2020) while Tk-GAL4^FruM^ cells were activated with an external LED (Fig. 3A). Upon LED stimulation, we observed localized increases in GCaMP6f fluorescence (Fig. 3B). The largest and most consistent change in fluorescence was observed in the *TkR86C^LexA^* neuronal processes that were near the SMP presynaptic sites of Tk-GAL4^FruM^ neurons (Fig. 3C). The activated domain extended posterior to the presynaptic area of Tk-GAL4^FruM^ neurons, while remaining clearly compartmentalized. We did not observe such an increase in calcium activity in areas overlapping with ring-adjacent postsynaptic projections (Fig. 3D). Although we occasionally observed fluorescence fluctuations in other areas of the brain (Fig. 3B_2_), this was weaker and less consistent than the activity in the SMP.

**Figure 3.**
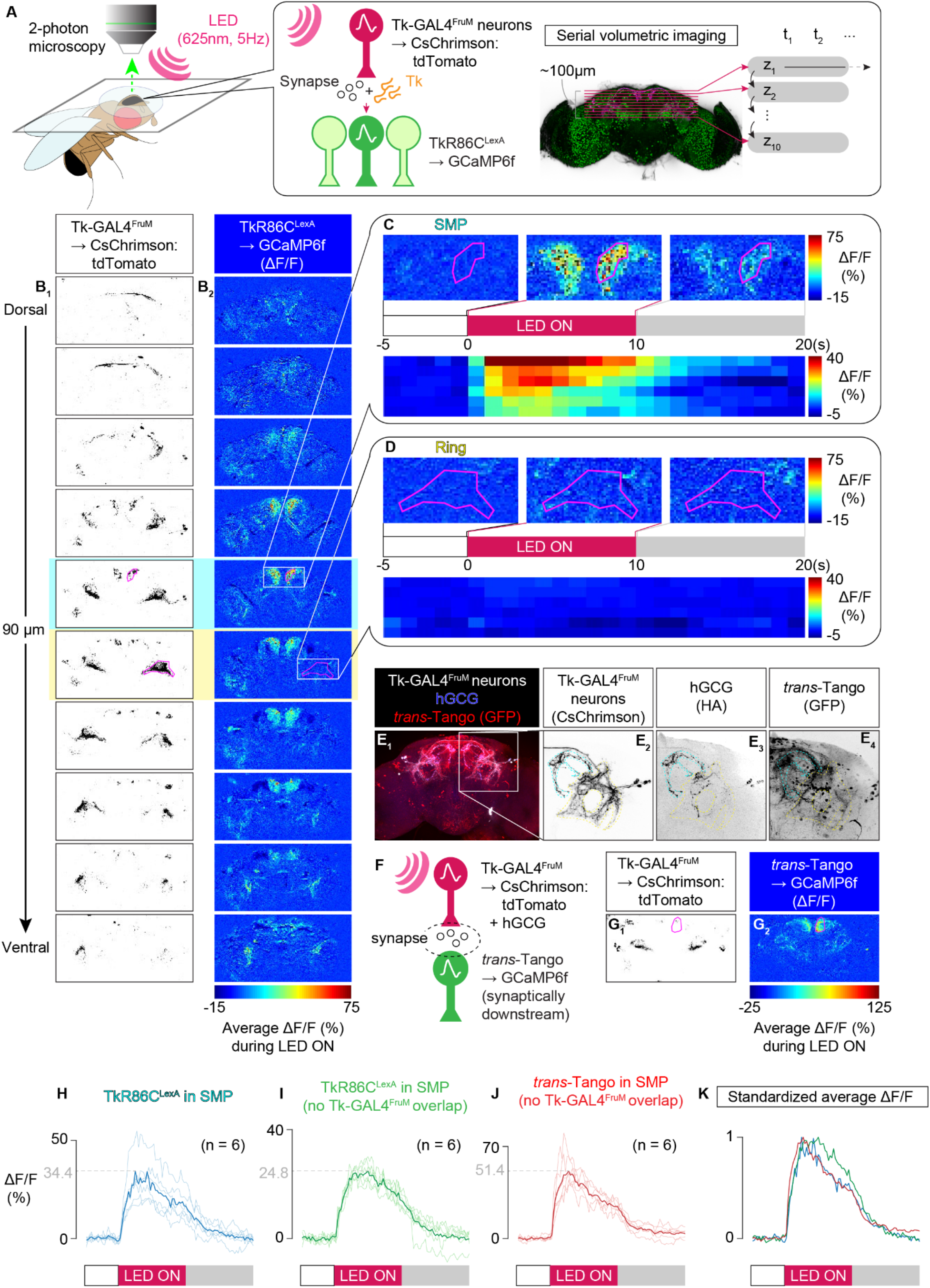
(previous page): *TkR86C^LexA^* neurons are likely postsynaptically activated by Tk-GAL4^FruM^ neurons. **A.** A schematic representation of the functional serial volume imaging. **B.** Images of CsChrimson:tdTomato fluorescence in Tk-GAL4^FruM^ neurons (**B1**) and the average fluorescence of GCaMP6f in *TkR86C^LexA^* neurons (**B2**) in a representative sample. The GCaMP6f signals are shown in pseudocolor as the relative increase in fluorescence (ΔF/F) during optogenetic activation of Tk-GAL4^FruM^ neurons. **C, D**. Pseudocolored GCaMP6f fluorescence changes (ΔF/F) in the vicinity of the projections of Tk-GAL4^FruM^ neurons to the SMP (**C**) and to the ring-adjacent region (**D**). Each series of three images shows the pixel-based ΔF/F taken from the sample shown in **B** and averaged over the time course before (left), during (middle), or after (right) stimulation. Below is the time course of fluorescence changes in similar regions of interest from six independent samples, binned into seconds and sorted by the average fluorescence change during stimulation, from most to least. **E**. A representative image of a male brain expressing CsChrimson:tdTomato (white in **E1**, **E2**) and membrane-tethered human glucagon (hGCG) (blue in **E1**, **E3**), the ligand of *trans*-Tango, in Tk-GAL4^FruM^ neurons. GFP expressed by the *trans*-Tango transgene is overlaid (red in **E1**, **E4**). **F**. A schematic representation of functional imaging of synaptically downstream partners of Tk-GAL4^FruM^ neurons using *trans*-Tango. **G**. Average fluorescence of CsChrimson:tdTomato in Tk-GAL4^FruM^ neurons (**G1**) and GCaMP6f (ΔF/F) in *trans-*Tango neurons (**G2**) from a slice posterior to the Tk-GAL4^FruM^ neuronal projection in the SMP in a representative brain during optogenetic activation of Tk-GAL4^FruM^ neurons. **H, I**. Fluorescence changes (ΔF/F) in *TkR86C^LexA^* neurons in the vicinity of (**H**) and in the area 10–20 µm posterior to (**I**) the Tk-GAL4^FruM^ neuronal projection in the SMP. **J**. Fluorescence changes (ΔF/F) in *trans-*Tango neurons in the area 10–20 µm posterior of Tk-GAL4^FruM^ neuronal projections in the SMP. Thick colored lines represent the average of the samples tested (thin lines). **K**. An overlay of the averages from **H-J**, normalized to the maximum fluorescence increase.

The fluorescence increase observed in the SMP began at the onset of LED stimulation and increased rapidly for about 2 s before starting to gradually decline even during the LED pulses (Fig. 3C). The fluorescence dropped when the LED was turned off, returning to the baseline in a few seconds in most cases. These spatial and temporal dynamics suggest that calcium activity in *TkR86C^LexA^* neurons is largely correlated with the activation of Tk-GAL4^FruM^ neurons. Importantly, these temporal dynamics were closely recapitulated when genetically defined, synaptically downstream neurons were accessed via the *trans*-Tango approach (Talay et al., 2017). Membrane-tethered human glucagon (hGCG) expressed in Tk-GAL4^FruM^ neurons drove expression of GCaMP6f in candidate synaptically downstream neurons (Fig. 3E_1-3_), which were many and scattered across the brain (Fig. 3E_1,4_). We then monitored LED stimulation-dependent calcium changes in these synaptically downstream neurons in response to optogenetic activation of Tk-GAL4^FruM^ neurons (Fig. 3F). Reflecting the rather widespread distribution of post-synaptic neurons, the fluorescent calcium activity was more widespread in *trans*-Tango samples than in brains expressing GCaMP6f under *TkR86C^LexA^* (Fig. S3A), and included activity in the ring-adjacent regions (Fig S3C). Part of the activity in the ring-adjacent area was generated by occasional GCaMP6f expression in Tk-GAL4^FruM^ neurons themselves (Fig. S3D, E), due to either lateral connectivity among Tk-GAL4^FruM^ neurons or self-labeling by *trans*-Tango (Fig S3F). Nonetheless, we consistently observed a fluorescence increase in the region posterior to (but not overlapping) the SMP projections of Tk-GAL4^FruM^ neurons (Fig. 3G, S3B). The activation patterns observed in the SMP were spatially and temporarily similar to the fluorescence dynamics observed in *TkR86C^LexA^* neurons (Fig 3H-K). Although we could not co-label *trans*- Tango neurons with *TkR86C^LexA^* due to lethality of the desired genotype, the functional imaging data support the notion that GCaMP6f signals in *TkR86C^LexA^* neurons result from direct postsynaptic connections with Tk-GAL4^FruM^ neurons.

### Cholinergic transmission is critical for the excitation of downstream *TkR86C^LexA^* neurons

The increase in intracellular calcium concentration in *TkR86C^LexA^* neurons with Tk-GAL4^FruM^ stimulation suggests that the overall impact of Tk-GAL4^FruM^ neuronal transmission is excitatory.

Consistent with this and with previous observations (Asahina et al., 2014), we found evidence that Tk-GAL4^FruM^ neurons co-express *choline acetyltransferase* (*ChAT*), a marker for excitatory cholinergic neurons (Fig. 4A), but not markers for glutamatergic (Fig. 4B) or GA-BAergic (Fig. 4C) neurons. Peptidergic ligands of *TkR86C* increase intracellular calcium concentration (Jiang et al., 2013; Poels et al., 2009), suggesting that the GCaMP6f signals we observed from *TkR86C^LexA^* neurons are a combination of cholinergic and tachykininergic transmission. To parse out the contribution of each of the two transmitter types, we first blocked cholinergic signaling with mecamylamine, an antagonist of the nicotinic acetylcholine receptor (Fig. 4D). The increase in GCaMP6f fluorescence in *TkR86C^LexA^* neurons triggered by optogenetic stimulation of Tk-GAL4^FruM^ neurons was severely reduced after bath application of mecamylamine, and could be partially rescued with a wash out (Fig. 4E_1-2_). By contrast, calcium signals remained largely unchanged when vehicle was added to the bath (Fig. 4F_1-2_). These data suggest that cholinergic signaling is a major contributor to the calcium activity observed in *TkR86C^LexA^* neurons upon Tk-GAL4^FruM^ neuronal activation.

**Figure 4:**
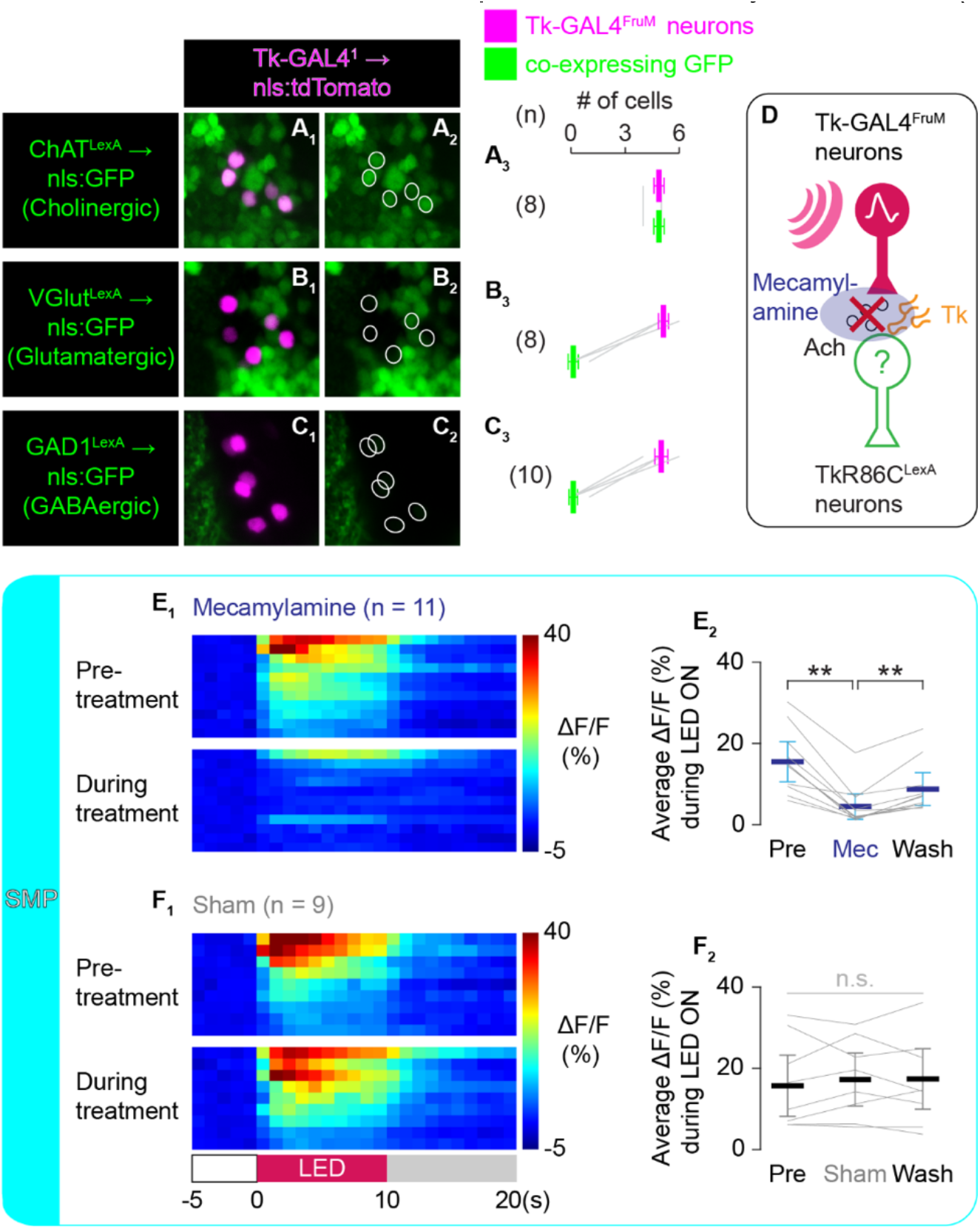
*TkR86C^LexA^* neurons receive cholinergic input from Tk-GAL4^FruM^ neurons. **A-C**. Overlap of nuclear-localizing tdTomato in Tk-GAL4^FruM^ neurons (magenta) and nuclear-localizing GFP in cholinergic (**A**), glutamatergic (**B**), and GABAergic (**C**) neurons (green). **D**. A schematic representation of the functional imaging experiment for **E** and **F** in which mecamylamine blocks cholinergic neurotransmission. **E, F**. Fluorescence change (ΔF/F) from *TkR86C^LexA^* neurons in the vicinity of the Tk-GAL4^FruM^ neuronal projection in the SMP with mecamylamine application (**E**) or sham treatment (**F**). Pseudocolored ΔF/F (**E1, F1**) represents fluorescence time courses for individual brains binned into seconds before (top) and during (bottom) treatment. Brain time courses are sorted by the average fluorescence change during stimulation, from most to least. The average ΔF/F from the same sample set before, during, and after treatment (**E2, F2**) is shown on the right. ** p < 0.01 by two-way ANOVA (p < 4.8 × 10^−8^) and post-hoc paired t-test. n.s. (in gray) p > 0.05 by two-way ANOVA. Thick lines and error bars in **A3**-**B3**, **E2**, and **F2** represent the average and 95% confidence intervals.

We reasoned that blocking synaptic transmission from *TkR86C^LexA^* neurons should prevent Tk-GAL4^FruM^ neurons from promoting aggression if these neurons are the major recipient of synaptic output from Tk-GAL4^FruM^ neurons. To test this possibility, we optogenetically activated Tk-GAL4^FruM^ neurons while blocking neurotransmission from *TkR86C^LexA^* neurons with the temperature-sensitive mutant protein of dynamin, Shibire^ts^ (Kitamoto, 2001). At a restrictive temperature of 32°C, where Shibire^ts^ is expected to block neurotransmission of *TkR86C^LexA^* neurons, optogenetic stimulation of Tk-GAL4^FruM^ neurons induced significantly fewer lunges in the mutant than in genetic controls (Fig. S4B,C). By contrast, at the permissive temperature of 22°C, the number of lunges during LED stimulation was comparable between the experimental and control genotypes (Fig. S4F,G), indicating that neurotransmission from *TkR86C^LexA^* neurons is necessary for Tk-GAL4^FruM^ neurons to promote aggression. Because *TkR86^LexA^* neurons are numerous in the nervous system, including in the ventral nerve cord (Fig. 2I), we cannot completely rule out a role for *TkR86^LexA^* neurons in general motor function. However, measures of other locomotor behaviors, such as distance traveled (Fig. S4D) and duration of orienting toward a target fly (Fig. S4E) (Wohl et al., 2020) during LED stimulation, were comparable in experimental and control genotypes, suggesting that blocking *TkR86C^LexA^* neuronal transmission does not impair basic motor function. Our data collectively argue that *TkR86C*-expressing neurons receive cholinergic synaptic inputs from Tk-GAL4^FruM^ neurons and are necessary for Tk-GAL4^FruM^-induced aggression.

### Tachykinin modulates excitatory postsynaptic responses in *TkR86C^LexA^* neurons

How does tachykinin modulate the cholinergic excitatory input from Tk-GAL4^FruM^ neurons onto *TkR86C^LexA^* neurons? To answer this question, we quantified the excitatory responses of *TkR86C^LexA^* neurons to optogenetic excitation of Tk-GAL4^FruM^ neurons while either eliminating (Fig. 5A, B) or over-expressing (Fig. 5C) *Tk*. As shown in Fig. 1, manipulating the amount of *Tk* changes how strongly Tk-GAL4^FruM^ neurons promote aggression upon optogenetic activation.

**Figure 5:**
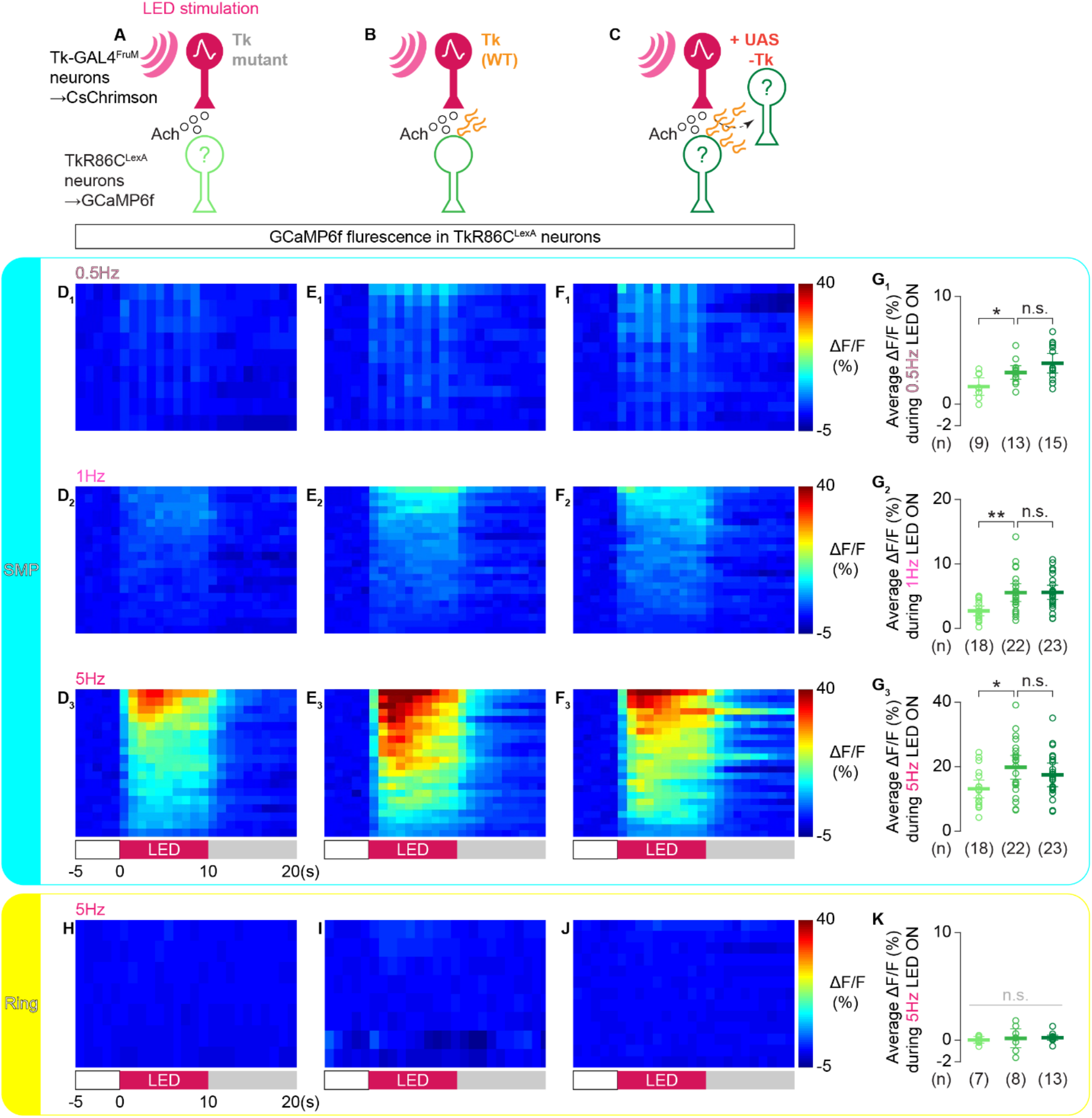
Tachykinin is necessary to maintain the wild-type intensity of excitatory synaptic transmission from Tk-GAL4^FruM^ neurons to *TkR86C^LexA^* neurons. **A-C.** Schematic representation of functional imaging experiments from *TkR86C^LexA^* neurons in *Tk* null mutants (**A**, data in **D, H**), *Tk* wild type (**B**, data in **E, I**), and *Tk* over-expression with a *UAS-Tk* transgene (**C**, data in **F, J**). **D-F**, **H-J**. Pseudocolored time course binned into seconds of GCaMP6f fluorescence changes (ΔF/F) in *TkR86C^LexA^* neurons, sorted by the average fluorescence change during stimulation, from the vicinity of the Tk-GAL4^FruM^ neuronal projection in the SMP (**D-F**) or in the ring-adjacent region (**H-J**). The SMP was imaged at three different LED frequencies as indicated above panels **D1-3**. **G, K.** Average ΔF/F in *TkR86C^LexA^* neurons during optogenetic stimulation of Tk-GAL4^FruM^ neurons, from the vicinity of the Tk-GAL4^FruM^ neuronal projection in the SMP (**G**) or in the ring-adjacent region (**K**). ** p < 0.01, * p < 0.05 by two-way ANOVA (**G1**: p < 2.7 × 10^−3^, **G2**: p < 6.9 × 10^−3^, **G3**: p = 0.017) and post-hoc paired t-test. n.s. (in gray, **K**) p > 0.05 by two-way ANOVA. Thick lines and error bars in **G1-3** and **K** represent the average and 95% confidence intervals. In **G** and **K**, the left data points are from *Tk* null mutants (light green), the middle data points are from *Tk* wild type (green), and the right data points are from animals with a *UAS-Tk* transgene (dark green), with n indicated at the bottom.

In *Tk* null mutants, the increase in GCaMP6f fluorescence in the SMP evoked by optogenetic stimulation of Tk-GAL4^FruM^ neurons was significantly attenuated compared with animals with the wild-type *Tk* locus (Fig. 5B), at multiple stimulation frequencies (Fig. 5D, E, G, S5A, B). The average increase in fluorescence (ΔF/F) was 30–50% lower in the *Tk* mutants than in wild type, which parallels the reduction in lunges induced by optogenetic stimulation of Tk-GAL4^FruM^ neurons under comparable LED power and frequencies (Fig. 1B, C, E). These data suggest that tachykinin is necessary for maintaining the strength of excitatory transmission between Tk-GAL4^FruM^ neurons and downstream *TkR86C^LexA^* neurons. The presence of responses in *TkR86C^LexA^* neurons in the *Tk* null background, albeit reduced, also suggests that acetylcholine alone can sustain some functional connectivity in the absence of tachykinin, reflecting the reduction but not elimination of aggression induced by Tk-GAL4^FruM^ excitation in *Tk* null mutants (Fig. 1B, E) (Asahina et al., 2014).

Interestingly, over-expression of tachykinin in Tk-GAL4^FruM^ neurons did not further increase GCaMP6f fluorescence in the SMP compared with the signals in animals with a wild-type *Tk* locus (Fig. 5E-G), even though the same genetic manipulation induced more lunges when Tk-GAL4^FruM^ neurons were activated at the same LED power and frequency (Fig. 1C-E). We did not observe any gross spatial changes in GCaMP6f signals from *TkR86C^LexA^* neurons when the *Tk* amounts were manipulated (Fig. S5B, C), including arbors near the ring-adjacent region of Tk-GAL4^FruM^ neurons (Fig. 5H-K). The absence of a difference in the response magnitude in the SMP may be due to the saturation of receptors in *TkR86C^LexA^* neurons. In fact, the level of receptor expression limits the efficacy of tachykininergic neuromodulation in olfactory and noticeptive circuits (Ignell et al., 2009; Im et al., 2015; Ko et al., 2015). However, over-expression of TkR86C in *TkR86C^LexA^* neurons did not further enhance aggression induced by the optogenetic activation of Tk-GAL4^FruM^ neurons (Fig. S5D, E). This suggests that the amount of tachykinin, rather than TkR86C receptors, is the limiting factor for the level of aggression. Moreover, optogenetic stimulation of Tk-GAL4^FruM^ neurons induced more lunges with *Tk* over-expression when the neurons also expressed GCaMP6f (Fig. S5F), excluding the possibility that GCaMP6f interferes with the aggression-promoting impact of *Tk* over-expression. These data collectively support the conclusion that excess tachykinin in *Tk-GAL4^1^* neurons does not change the dynamics of the circuit that involves *TkR86C^LexA^* neurons, even though it both quantitatively and qualitatively enhances aggression induced by the optogenetic activation of Tk-GAL4^FruM^ neurons.

### Tachykinin overexpression in Tk-GAL4^FruM^ neurons recruits *TkR99D*-expressing neurons

The absence of a noticeable difference in *TkR86C^LexA^* calcium signals with tachykinin over-expression suggests that these are not the only neural correlates of enhanced aggression induced by activation of Tk-GAL4^FruM^ neurons (Fig. 1C-H). We asked whether another tachykinin receptor, *TkR99D* (Birse et al., 2006), plays a role in defining a parallel behaviorally relevant circuit. Although not required for aggression induced by the activation of Tk-GAL4^FruM^ neurons (Asahina et al., 2014), TkR99D receptor proteins may detect over-expressed tachykinin from Tk-GAL4^FruM^ neurons (which can increase the local concentration of tachykinin), perhaps without direct synaptic connection, given the higher affinity of this receptor to tachykinin than TkR86C (Birse et al., 2006; Jiang et al., 2013; Poels et al., 2009). To address this possibility, we created a LexA knock-in allele of *TkR99D* by the same strategy used for *TkR86C^LexA^* (Fig. S6A, B). Like *TkR86C^LexA^*, *TkR99D^LexA^* labeled many neurons throughout the brain (Fig. 6A), but not TkGAL4^FruM^ neurons themselves (Fig. 6B_1-2_, C). In contrast to *TkR86C^LexA^* neurons, the overlap of *TkR99D^LexA^* neurons near the presynaptic projections of Tk-GAL4^FruM^ neurons in the SMP was comparable to that in the postsynaptic regions (Fig. 6B_3-4_, D).

**Figure 6:**
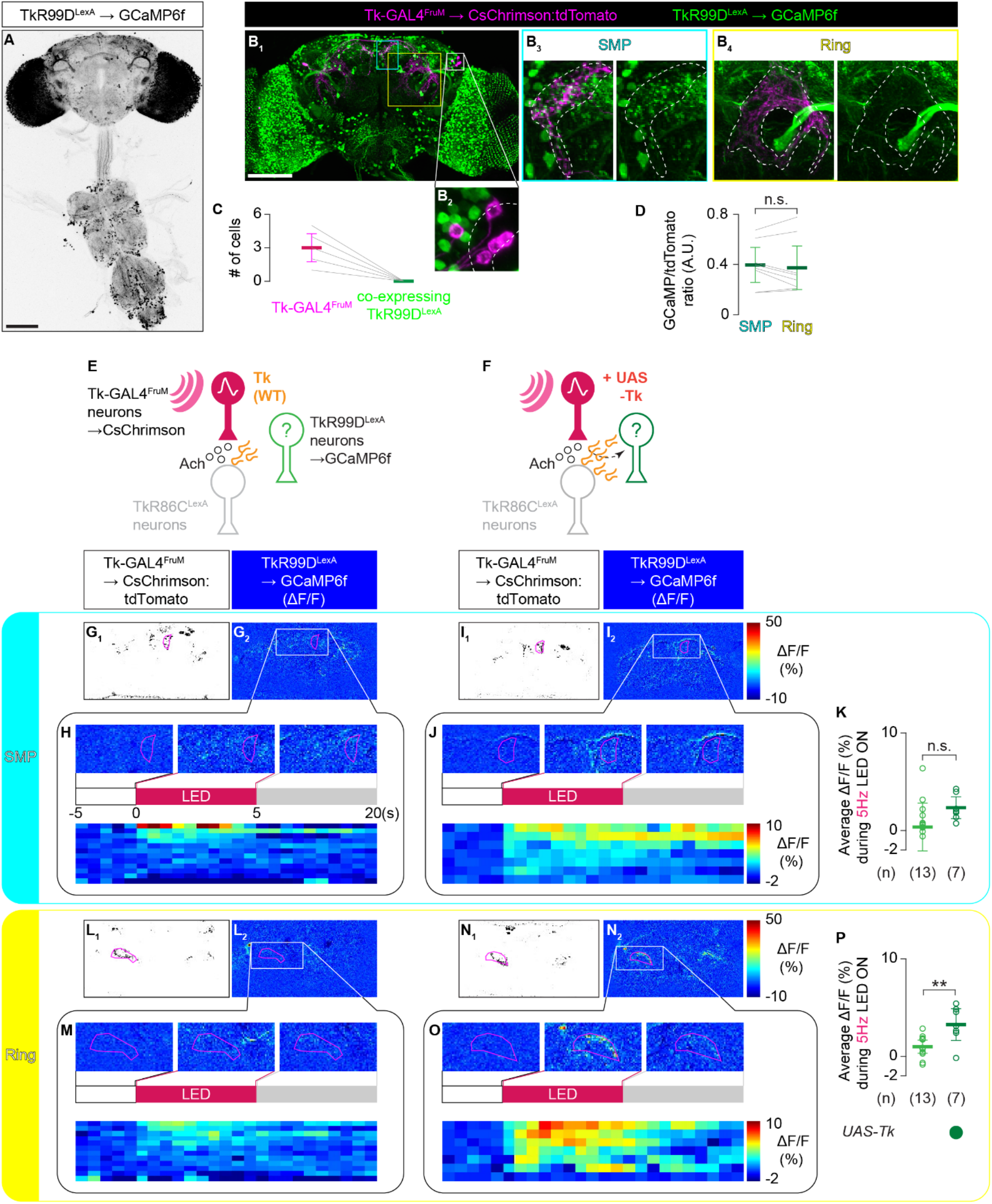
*TkR99D^LexA^* neurons are consistently activated by Tk-GAL4^FruM^ neurons when tachykinins are over-expressed. **A.** Representative expression pattern of GCaMP6f driven by *TkR99D^LexA^* in the nervous system, visualized by immunohistochemistry. Scale bar: 100 µm. **B.** Representative image of a male brain expressing GCaMP6f driven by *TkR99D^LexA^* (green) and CsChrimson:tdTomato under intersectional control of *Tk-GAL4^1^* and *fru^FLP^* (magenta). Images near the Tk-GAL4^FruM^ cell bodies (**B2**) and their projections (broken white lines) in the SMP (**B3**) and the ring (**B4**) regions are magnified. Scale bar: 100 µm. **C.** *TkR99D^LexA^* does not label Tk-GAL4^FruM^ neurons (n = 6). **D.** Boxplot of GCaMP6f immunohistochemical signals in the SMP and ring-adjacent regions defined by Tk-GAL4^FruM^ neurons, relative to their tdTomato immunohistochemical signals. n.s. p > 0.05 by paired t-test. **E, F**. Schematic representations of functional imaging experiments from *TkR99D^LexA^* neurons in *Tk* wild type (**E**, for data in **G**, **H**, **L**, **M**) and in the presence of a *UAS-Tk* transgene (**F**, for data in **I**, **J**, **N**, **O**). **G, I, L, N.** Images of CsChrimson:tdTomato fluorescence in Tk-GAL4^FruM^ neurons (**G1**, **I1**, **L1**, **N1**) and average GCaMP6f signals in *TkR99D^LexA^* neurons (**G2**, **I2**, **L2**, **N2**) during optogenetic activation of Tk-GAL4^FruM^ neurons, from representative sample in the optical slices that include the Tk-GAL4^FruM^ neuronal projections in the SMP (**G, I**) or in the ring-adjacent region (**L, N**). Pseudocolor GCaMP6f signals are the relative increase in fluorescence (ΔF/F) as indicated on the right. **H, J, M, O**. Pseudocolored GCaMP6f fluorescence changes (ΔF/F) from the vicinity of the projections from Tk-GAL4^FruM^ neurons to the SMP (**H**, **J**) or the ring-adjacent region (**M**, **O**), represented as the spatiotemporal average before (left), during (middle), and after (right) Tk-GAL4^FruM^ stimulation of the sample shown above (top) and as a time course, binned into seconds, from 13 (**H**, **M**) or 7 (**J**, **O**) independent samples sorted by average fluorescence change during stimulation, from most to least (bottom). **K, P.** Average ΔF/F in *TkR99D^LexA^* neurons during optogenetic stimulation of Tk-GAL4^FruM^ neurons, from the vicinity of the Tk-GAL4^FruM^ neuronal projection in the SMP (**K**) or in the ring-adjacent region (**P**). In **K** and **P**, left data points are from *Tk* wild type and right data points are from animals with a *UAS-Tk* transgene, with n indicated at the bottom. ** p < 0.01, n.s. p > 0.05 by t-test.

We next asked whether any *TkR99D^LexA^* neurons are functionally downstream of Tk-GAL4^FruM^ neurons. We expressed GCaMP6f under the control of *TkR99D^LexA^* while expressing CsChrimson in Tk-GAL4^FruM^ neurons (Fig. 6E), and monitored fluorescence intensity in response to optogenetic stimulation of Tk-GAL4^FruM^ neurons. We did not observe consistent fluorescence fluctuations near the innervation from Tk-GAL4^FruM^ neurons, either in the SMP (Fig. 6G, H) or in the ring-adjacent region (Fig. 6L, M, S6C). We noticed that GCaMP6f intensity often increased after LED stimulation in the protocerebral bridge (Fig. S6C, D), where Tk-GAL4^FruM^ neurons do not project. This neural structure is known to respond to visual stimuli in both *Drosophila* (Weir and Dickinson, 2015) and other insect species (Heinze and Homberg, 2007; Homberg et al., 2011; Pegel et al., 2019; Phillips-Portillo, 2012). Therefore, direct activation of this visual circuit by the LED light may have led to the observed calcium response.

Interestingly, when *Tk* was over-expressed in Tk-GAL4^FruM^ neurons (Fig. 6F), optogenetic activation elevated GCaMP6f fluorescence near Tk-GAL4^FruM^ neurons (Fig. 6I, J, N, O, S6D). The fluorescence increase near the ring-adjacent region of Tk-GAL4^FruM^ neurons was significantly higher than in animals with wild-type *Tk* loci (Fig. 6N-P), while the signal in the SMP remained comparable (Fig 6K). These newly recruited *TkR99D^LexA^* neurons are distinct from the *TkR86C^LexA^* neurons that are synaptically downstream of Tk-GAL4^FruM^ neurons, since *TkR86C^LexA^* neurons near the ring-adjacent region are not recruited by optogenetic activation of Tk-GAL4^FruM^ neurons that over-express *Tk*. This suggests that tachykinins released from Tk-GAL4^FruM^ neurons can modulate two distinct circuits depending on the available amount of *Tk*. Our data support the idea that *Tk* overexpression in Tk-GAL4^FruM^ neurons potentiates their aggression-promoting capability by recruiting an additional population of neurons that receive tachykinin via *TkR99D*.

## DISCUSSION

Although neuropeptides modulate a wide range of behaviors, the cellular and genetic basis of this modulation has remained elusive. Using functional imaging, we found that tachykinin released from Tk-GAL4^FruM^ neurons can modulate two distinct circuits (Fig. 7). One is likely a direct postsynaptic target that expresses *TkR86C*. These neurons are necessary for Tk-GAL4^FruM^ neurons to promote aggression, with tachykinin modulating the excitatory response triggered by the co-transmitter acetylcholine. The other circuit is labeled by *TkR99D*. While largely non-responsive to Tk-GAL4^FruM^ neuronal stimulation, a higher level of tachykinin in Tk-GAL4^FruM^ neurons recruits these neurons; this may account for both qualitative and quantitative enhancement of aggressive behaviors when tachykinin is over-expressed in Tk-GAL4^FruM^ neurons. Our results uncover a mechanism by which neuropeptides engage multiple neural circuits labeled by distinct neuropeptide receptors to control behavior intensity.

**Figure 7:**
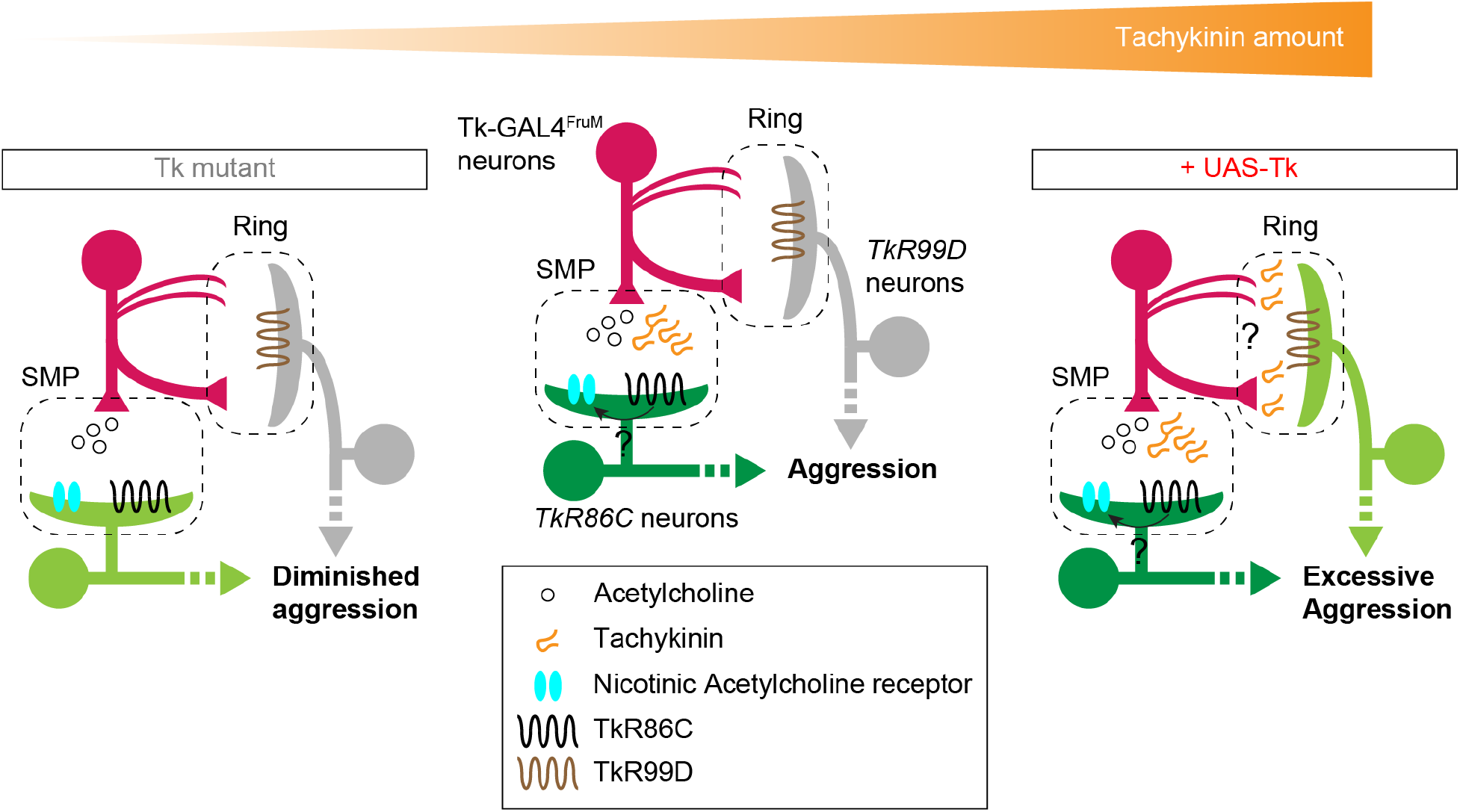
A model of tachykinin-mediated circuit reconfiguration. Tk-GAL4^FruM^ neurons make cholinergic synaptic connections in the SMP with *TkR86C*-expressing downstream neurons, which mediate the aggression promoted by Tk-GAL4^FruM^ neurons (middle). Tachykinin potentiates the excitatory transmission through an unknown mechanism. In the absence of tachykinin (left), the *TkR86C*-expressing downstream population is not as effectively excited by Tk-GAL4^FruM^ neurons, resulting in a diminished level of aggression. Over-expression of tachykinin in Tk-GAL4^FruM^ neurons recruits *TkR99D*-expressing neurons that project to the ring-adjacent region. Although Tk-GAL4^FruM^ neurons send mainly postsynaptic arbors to the ring area, axon termini from the contralateral side also reach there. Tachykinin from either structure excites *TkR99D*-expressing neurons, which we speculate may contribute to excessive aggression.

### Division of labor by different peptidergic receptors

A single neuropeptide species is often recognized by multiple receptors (Griebel and Holsboer, 2012; Nässel and Winther, 2010). Different receptors are often expressed in separate neuronal populations, suggesting that they delineate neural circuits that are distinct from one another. Although we are currently unable to visualize the overlap of *TkR86C^LexA^* neurons and *TkR99D^LexA^* neurons directly, our data support the idea that *TkR86C^LexA^* neurons and *TkR99D^LexA^* neurons that are activated by Tk-GAL4^FruM^ neurons are non-overlapping populations. First, they are spatially segregated. Optogenetic activation of Tk-GAL4^FruM^ neurons excites *TkR86C^LexA^* neurons located almost exclusively near the axon termini of Tk-GAL4^FruM^ neurons in the SMP, whereas the same manipulation excites *TkR99D^LexA^* neurons that have processes near the dendritic arbors of Tk-GAL4^FruM^ neurons in the ring-adjacent region. Second, *TkR86C^LexA^* neurons can be excited by optogenetic activation of Tk-GAL4^FruM^ neurons even in the absence of tachykinin peptides, whereas *TkR99D^LexA^* neurons are reliably excited only when *Tk* is over-expressed in Tk-GAL4^FruM^ neurons. While it remains possible that the two tachykinin receptors are co-expressed in some neurons, *TkR86C* and *TkR99D* are used by separate neuronal populations in circuits modulated by tachykinins released from Tk-GAL4^FruM^ neurons. Our data depict a model in which neuropeptides from a single population of neurons sculpt the activity in two separate downstream targets defined by different receptors (Fig. 7).

A unique feature of peptidergic neuromodulation is the diversity of neuronal targets (Nässel, 2009; Nusbaum et al., 2017; van den Pol, 2012). Peptidergic neuromodulation can (1) act upon synaptically downstream partners to potentiate or attenuate the impact of classical neurotransmitters, (2) mediate dendro-dendritic or retrograde interactions through release from presynaptic areas, (3) cause auto-modulation by activating receptors expressed by the peptide-releasing neurons themselves, and (4) exert long-range effects as neurohormones. Our brain-wide functional imaging revealed restricted activity patterns in response to optogenetic stimulation of Tk-GAL4^FruM^ neurons, suggesting tachykinins in this context mainly act locally. The absence of *TkR86C^LexA^* or *TkR99D^LexA^* expression in Tk-GAL4^FruM^ neurons excludes auto- or axoaxonal modulation of Tk-GAL4^FruM^ neurons. The spatiotemporal similarity of the activity patterns in genetically identified postsynaptic neurons and *TkR86C^LexA^* neurons suggest that *TkR86C* mediates postsynaptic enhancement of cholinergic neurotransmission. The fact that an acetylcholine receptor antagonist almost completely blocks the Tk-GAL4^FruM^ neuron-induced activity in *TkR86C^LexA^* neurons further supports this conclusion. The relationship between *TkR99D^LexA^* neurons and Tk-GAL4^FruM^ neurons remains unclear. Since *TkR99D^LexA^* neurons are activated in proximity to dendritic areas of Tk-GAL4^FruM^ neurons, it is possible that they receive tachykinin released from the dendrites of Tk-GAL4^FruM^ neurons. In this scenario, *TkR99D* could mediate either parallel (dendro-dendritic) or retrograde modulation by tachykinin. On the other hand, ring-adjacent postsynaptic neurons that express *TkR99D* may be activated by the contralateral projection of Tk-GAL4^FruM^ neurons. Identification of specific receptor-expressing target neurons will clarify these possibilities. Characterization of the synaptic structure of male-specific neurons such as Tk-GAL4^FruM^ neurons remains a challenge because electron-microscopy-based wiring diagrams are only available for the female fly brain (Scheffer et al., 2020; Zheng et al., 2018). Our multiple attempts to identify candidate synaptic partners using photo-activatable GFP (Aso et al., 2014; Datta et al., 2008; Ruta et al., 2010) have been unsuccessful (data not shown). *Trans*-Tango expressed in Tk-GAL4^FruM^ neurons labeled hundreds of cells across the brain (sometimes including Tk-GAL4^FruM^ neurons themselves), preventing us from characterizing the neuroanatomy of each of them.

Nonetheless, our data outline how neuropeptides from a single group of neurons can functionally reconfigure different receptor-expressing neurons in a peptide dose-dependent manner. The existence of multiple receptors is known to be important for diversifying neuromodulator targets. In vertebrates, D1 and D2 dopamine receptors label largely non-overlapping sub-populations of medium spiny neurons (Gerfen et al., 1990; Gong et al., 2007), which play complementary roles in motion control (Geddes et al., 2018; Jin et al., 2014). In *Drosophila*, different dopamine receptors play distinct roles in both innate (Sayin et al., 2019; Zhang et al., 2016) and learned (Handler et al., 2019) behaviors, at least in part by activating different downstream signaling cascades (Handler et al., 2019). As for neuropeptides, diuretic hormone 44 (Dh44) released from the glucose-sensing neurons in the central brain of *Drosophila* acts on two distinct downstream target neurons labeled by expression of two different receptors, *Dh44-R1* (in downstream neurons) and *Dh44-R2* (in gut cells) (Dus et al., 2015). These two cell types coordinate starvation-induced behavioral and physiological changes. Collectively, these examples depict a motif whereby multiple receptors of a neuromodulator define functionally distinct downstream circuits. Our results indicate that different downstream targets of aggression-promoting Tk-GAL4^FruM^ neurons are recruited depending on the peptide level from a single cluster of neurons, contributing to distinct aspects of behavioral escalation.

### Modes of tachykininergic neuromodulation

Tachykinins constitute an evolutionarily conserved family of neuropeptides (Nässel et al., 2019; Severini et al., 2002). It is intriguing that tachykinins are known to control aggressive behaviors in several mammalian species (Katsouni et al., 2009; Zelikowsky et al., 2018). While vertebrate tachykinins (such as substance P) are considered excitatory neuropeptides (Jan and Jan, 1982; Phillis and Limacher, 1974), *Drosophila* tachykinin is known to pre-synaptically inhibit olfactory sensory neurons in the antennal lobe (Ignell et al., 2009; Ko et al., 2015). Our study demonstrates that *Drosophila* tachykinin also serves as an excitatory neuromodulator. In a heterologous cell expression system, both *TkR86C* and *TkR99D* receptors generate an increase in calcium concentration upon application of tachykinin (Birse et al., 2006; Jiang et al., 2013; Johnson et al., 2003; Poels et al., 2009), consistent with our functional imaging data. Molecular mechanisms that support opposing modulatory effects of tachykinins in olfactory and aggression-controlling circuits is an important topic for further investigation. *TkR99D* is involved in both excitatory and inhibitory modulation, suggesting that it may be capable of coupling with both excitatory and inhibitory G-proteins in different neuronal populations. Alternatively, different tachykinin neuropeptides may have different pharmacological impacts on these receptors. It is known that all six mature peptides (DTK1-DTK6) generated from the tachykinin prepropeptide activate *TkR99D* (Birse et al., 2006; Jiang et al., 2013), whereas *TkR86C* can be activated only by relatively high concentration of DTK6 (Jiang et al., 2013; Poels et al., 2009). In fact, a preferred ligand of *TkR86C* is another neuropeptide, natalisin (Jiang et al., 2013). Although it remains unknown whether Tk-GAL4^FruM^ neurons release natalisin, tachykininergic neurons in different microcircuits may release distinct mixes of mature tachykinins, which could modulate receptor-expressing neurons in each of these microcircuits in physiologically distinct manners. Lastly, tachykinin receptors can engage multiple intracellular signaling cascades. In the current study, we monitored Ca^2+^ as a read-out of neuromodulation and excitatory neurotransmission. Visualization of other signaling components such as cAMP (Shafer et al., 2008) would provide a more comprehensive understanding of the cellular impact of tachykininergic modulation. By resolving these possibilities, we will not only advance our understanding of peptidergic neuromodulation, but also help more accurately predict the physiological and behavioral effects of pharmacological substances that are designed to target specific receptor-expressing neurons (Griebel and Holsboer, 2012; Holmes et al., 2003).

A neuromodulator can affect circuits and behavior in a functionally distinct way from a co-expressed neurotransmitter, as shown both in flies (Sherer et al., 2020) and in mice (Chen et al., 2019; Zell et al., 2020). Because neuromodulators (especially neuropeptides) may communicate with receptor-expressing neurons extra-synaptically, the connectome by itself may not fully reveal the physiologically and behaviorally relevant functional relationships among neurons. The expression profiles of neuromodulator receptors (coined the “chemoconnectome” (Deng et al., 2019)) in these aggression-controlling neuromodulatory cells can provide an insight into the functional connectivity among them.

### Neural mechanism of peptidergic modulation of aggressive behavior

In addition to tachykinins, many neuropeptides are known to regulate aggressive behavior, in flies as well as in vertebrates (Agrawal et al., 2020; Albers, 2012; Asahina, 2017; Katsouni et al., 2009; Wu et al., 2020). The dynamic nature of peptidergic neuromodulation is particularly beneficial for aggressive behavior, which must be adjusted both qualitatively and quantitatively according to resource availability, previous fighting and other social experiences, and the perceived fighting capacity of an opponent (Maynard Smith, 1982). In this study, we showed that tachykinins modulate a *TkR86C*-expressing circuit to quantitatively control the intensity of aggression triggered by the activation of Tk-GAL4^FruM^ neurons. The presence of tachykinin is necessary within this circuit to maintain the strength of excitatory synaptic transmission, and that transmission is necessary for the production of lunges. On the other hand, a *TkR99D*-expressing circuit likely mediates qualitative, as well as quantitative, enhancement of aggression caused by the over-expression of tachykinins in Tk-GAL4^FruM^ neurons. Under this genetic manipulation, activation of Tk-GAL4^FruM^ neurons drives male flies to attack a moving small magnet (Asahina et al., 2014) and a female fly, behaviors which are seldom observed in normal males (Fernández et al., 2010; Monyak et al., 2021). Our data suggest that the limiting factor in peptidergic transmission strength is the amount of released peptide, and not the level of receptor expression like in the adult olfactory circuit (Ignell et al., 2009; Ko et al., 2015) or larval noticeptive circuit (Im et al., 2015). This means that dynamic regulation of peptide amount released from Tk-GAL4^FruM^ neurons may instruct both the intensity and target of aggressive actions. Knowledge of the cellular mechanisms that determine the release quantity of tachykinins from Tk-GAL4^FruM^ neurons will be critical for understanding how Tk-GAL4^FruM^ neurons integrate the internal and external factors that are known to affect the level of fly aggression.

How tachykininergic systems interface with other peptidergic systems, such as NPF (Dierick and Greenspan, 2007) and Drosulfakinin (Agrawal et al., 2020; Wu et al., 2020), or biogenic amine neuromodulators (Alekseyenko et al., 2013, 2014, 2019; Andrews et al., 2014; Certel et al., 2010; Dierick and Greenspan, 2007; Hoyer et al., 2008; Watanabe et al., 2017; Zhou et al., 2008), remains an important question to be resolved. Identification of the context in which each population is engaged is an important step in delineating the contributions of each neuro-modulator-releasing neuron group. It is possible that each neuromodulator reflects a specific internal or external condition that helps the animal weigh the costs and benefits of fighting. In the case of the tachykininergic system, characterization of the neural inputs into Tk-GAL4^FruM^ neurons and determinants of tachykinin release amount will help us understand which aspects of strategic decision-making are mediated by this population of neurons, and how tachykinins serve as a molecular actuator of the consequential behavioral choices.

## ACKNOWLEDGEMENTS

We thank Yoshi Aso, Jinyang Liu, and Steven Sawtelle (Janelia Research Campus) for sharing the FlyBowl acquisition software; David Tsu and Eric De la Parra for their assistance in fly maintenance and behavioral assays. M.W. was supported by the Mary K. Chapman Foundation and Rose Hills Foundation. This work has been supported by NIH NIDCD R01 DC015577 to K.A. K.A. is a recipient of the Helen McLoraine Development Chair of Neurobiology at the Salk Institute.

## AUTHOR CONTRIBUTIONS

Conceptualization, K.A.; Methodology, K.A., M.W.; Software, M.W.; Formal Analysis, K.A., M.W.; Investigation, K.A., M.W.; Resources, K.A., M.W.; Data Curation, K.A., M.W.; Writing – Original Draft, K.A., M.W.; Writing – Review & Editing, K.A., M.W.; Supervision, K.A.; Funding Acquisition, K.A.

## MATERIALS AND METHODS

### Fly Strains

See Table 1 for the complete genotypes of *Drosophila* strains used in each figure panel. *Tk-GAL4^1^* (RRID:BDSC_51975), *Otd-nls:FLPo* (in attP40), *ΔTk^1^*, *10XUAS-Tk* were previously described in (Asahina et al., 2014). *20XUAS>myr:TopHAT2>CsChrimson:tdTomato* (in VK00022 and VK00005) (Duistermars et al., 2018; Watanabe et al., 2017), *13XLexAop2-IVS-Syn21-GCaMP6f* (codon-optimized)*-p10* (in su(Hw) attP5 and su(Hw)attP1) and *13XLexAop2-IVS-syn21-shibire^ts^-p10* (in VK0005) (Pfeiffer et al., 2012) were created by Barret Pfeiffer and kindly shared by David Anderson (California Institute of Technology) and Gerald Rubin (HHMI Janelia Research Campus) *fru^FLP^* (RRID:BDSC_66870) (Yu et al., 2010) was a gift from Barry Dickson (HHMI Janelia Research Campus). *pJFRC118-10XUAS-TLN:mCherry* (DenMark) (in attP40) and *pJFRC67-3XUAS-IVS-Syt:GFP* (in Su(Hw)attP1) (Seelig and Jayaraman, 2013) were gift from David Anderson (California Institute of Technology). *trans*-Tango (in attP40) (RRID:BDSC_77123) (Talay et al., 2017) and *QUAS-mCD8:GFP* (Potter et al., 2010) were a gift from Mustafa Talay and Gilad Barnea (Brown University). *Tubulin-FRT-GAL80-FRT-stop* (Gordon and Scott, 2009) was a gift from Kristin Scott (UC Berkeley). *hs-Cre* (Siegal and Hartl, 1996) (RRID:BDSC_851), *vasa-Cas9* (Gratz et al., 2014) (RRID:BDSC_51323), *VGlutLexA:QFAD.2* (RRID:BDSC_60314), *ChAT-LexA:QFAD.0* (RRID:BDSC_60319), and *Gad1-LexA:QFAD.2* (RRID:BDSC_60324) (Diao et al., 2015) flies were obtained from Bloomington Drosophila Resource Center at the University of Indiana.

### Creation of knock-in strains

*Takr86C^LexA^* and *Takr99D^LexA^* knock-in alleles were created using CRISPR/Cas9-mediated genome editing (Gratz et al., 2014). For both *TkR86C* and *TkR99D*, we first identified a pair of 21-nucleotides guide RNA (gRNA) sequences, using flyCRISPR Target Finder (http://target-finder.flycrispr.neuro.brown.edu/), that are expected to delete the segment between the start codon and 3’ end of the first coding sequence-containing exon. The gRNA sequences are listed below (PAM sequences are underlined):

*TkR86C* gRNA #1: GCAGTCTGTAATCAGGATAG <u>AGG</u>
*TkR86C* gRNA #2: GTACTTCCTGCCCACTCACT <u>TGG</u>
*TkR99D* gRNA #1: GAAGTCACTGCGATTCTCCA <u>TGG</u>
*TkR99D* gRNA #2: GTCATAATTAGGCATGCCGG <u>CGG</u>

Two gRNA sequences for each gene were incorporated into the tandem gRNA expression vector pCFD4 following the protocol described in (Port et al., 2014). We call this plasmid a gRNA plasmid. In parallel, we also created a donor plasmid for each gene, using pHD-DsRed (Addgene cat#51434) (Gratz et al., 2014) as a backbone. The donor plasmid contains the coding sequence of LexA:p65 (Pfeiffer et al., 2010) in frame with the start codon of *TkR86C* or *TkR99D*. These coding sequences are sandwiched by the 5’ UTR and the sequence immediately downstream of the start codon of *TkR86C* or *TkR99D*. The floxed 1,225bp 3XP3-DsRed-SV40 marker gene was inserted in the orientation opposite to the targeted gene in the intron region of the 3’ arm (1,294 – 70 bp downstream of the 3’ end of 1^st^ exon of *TkR86C*, and 1,293 – 69 bp downstream of the 3’ end of 2^nd^ exon of *TkR99D*). The start codon of *TkR86C* or *TkR99D* in the donor plasmid was changed to the amber stop codon (TAG). Also, the PAM motifs of the gRNA sequences on both arms within the donor plasmid were mutated so as to avoid secondary cleavage by Cas9 proteins. DNA fragments for both 5’- and 3’-homologous arms were amplified using PrimeSTAR GXL DNA polymerase (TaKaRa Clontech cat# R050) from the genome DNA of Canton-S wild-type strain of *D. melanogaster*, which contained several point mutations and small indels compared to the standard *Drosophila* genome sequence. The 5’ arm and LexA:p65 coding sequence was assembled from 2 fragments from PCR-amplified *Drosophila* genome and a LexA:p65 coding sequence using NEBuilder HiFi DNA Assembly Cloning Kit (New England Biolabs, cat#E5520), and inserted into XhoI-SpeI sites of the pHD-DsRed plasmid. The 3’ arm was subsequently inserted into NdeI-EcoRI sites of the intermediate plasmid using the same kit. The sequence of the plasmids that is expected to be incorporated into the fly genome was verified by Sanger sequencing.

The appropriate combination of gRNA and donor plasmids was mixed and injected into embryos of *vasa-Cas9* strain (Bloomington #51323) by BestGene Inc. G1 adults (offspring of injected (G0) animals) were screened for the presence of DsRed expression in the compound eyes, followed by PCR screening. The Southern blotting was used to verify the correct integration of the donor element (see below). After backcrossing the knock-in alleles in a Canton-S background for 6 generations, the 3XP3-DsRed marker gene was removed by using *Cre* recombinase. Specifically, flies containing the knock-in allele crossed to flies that express the *hs-Cre* transgene (Siegal and Hartl, 1996). This transgene induced efficient excision of the floxed marker gene under the standard rearing temperature of 25°C, as *hs-Cre* was previously reported to be active without heat shock (Hampel et al., 2011; Siegal and Hartl, 1996). The offspring were screened for the loss of DsRed expression in the eyes.

### Creation of transgenic strains

The *15XQUAS-GCaMP6f* (in su(Hw)attP5 and su(Hw)attP1) transgenic strains were created in the following steps. First, a DNA fragment that contains IVS-Syn21-GCaMP6f (codon optimized)-p10 elements was amplified from the genomic DNA of the transgenic strain that carries *13XLexAop2-IVS-Syn21-GCaMP6f-p10* (in su(Hw)attP1) by PCR (Phusion Green, ThermoFisher Scientific cat# F534). This fragment was subcloned into pCR Blunt II TOPO vector using the Zero Blunt TOPO kit (ThermoFisher Scientific cat# K287540). In parallel, a modified version of the plasmid pJFRC164-21XUAS-KDRT>-dSTOP-KDRT>-myr::RFP (Addgene #32141), in which the 21XUAS element was replaced with a 13XLexAop2 element, was digested with XhoI and EcoRI. The IVS-Syn21-GCaMP6f-p10 element in the pCR Blunt II TOPO vector was amplified with overhang sequences, and ligated into the digested backbone of the modified pJFRC164 using the In-Fusion HD Cloning Kit (TaKaRa Clontech cat#639648) to create the plasmid 13XLexAop2-KDRT>-dSTOP-KDRT>-IVS-Syn21-GCaMP6f-p10 intermediate plasmid (named pMW02). Next, the 15XQUAS sequence from the plasmid pBAC-ECFP15XQUAS-TATA-mCD8:GFP-SV40 (Addgene #104878) was amplified by PCR, which was subsequently used to replace the LexAop2 sequence of pMW02, which was excised by HindIII and AatII, using the In-Fusion HD Cloning Kit. The resulting plasmid, 15XQUAS-KDRT>-dSTOP-KDRT>- IVS-Syn21-GCaMP6f-p10, was then digested with AatII and NotI to remove the KDRT cassette, which was replaced by a Hsp70-IVS fragment excised by AatII and NotI from the plasmid pJFRC28-10XUAS-IVS-GFP-p10 (Addgene #36431). The sequence of the final product (15XQUAS-Hsp70-IVS-Syn21-GCaMP6f (codon-optimized)-p10, shorthanded as 15XQUAS-GCaMP6f) was verified before being integrated into target attP sites via phiC31-mediated site-specific transformation (BestGene Inc.,).

The *LexAop2-TkR86C* transgenic element was created by replacing the myr:GFP coding sequence of the plasmid pJFRC19-13XLexAop2-IVS-myr::GFP (Addgene #26224) with the coding sequence of *TkR86C*. Specifically, a DNA fragment of the *TkR86C* coding region was amplified from cDNA from the Canton-S wild type strain by PCR (PrimeSTAR GXL, TaKaRa Clontech) with primers that had NotI and XbaI sites at 5’ and 3’ ends, respectively. The fragment was subcloned into the pCR Blunt II TOPO vector. pJFRC19 plasmids and *TkR86C*-containing vector plasmids were digested with NotI and XbaI. The pJFRC19 backbone and *TkR86C* fragments were ligated using Roche Rapid DNA Ligation Kit (Millipore Sigma cat#11635379001). The recovered *TkR86C* coding sequences (isoform B, 1,665bp) have 3 base substitutions, including 1 non-synonymous mutation (T425I), compared to the NCBI reference sequence NP_001097741.1. The HA-tagged version was created by adding the 135bp that contains a 3X repeat of the hemagglutinin sequence at the C-terminus of the *TkR86C* coding sequence. The coding region was fully sequenced before transformation.

The sequences of the DNA constructs used in this study are shown in Table 2.

### Southern Blotting

Two hundred adult flies per genotype were homogenized in 800 µL of TE buffer (Tris/HCl (pH 9), 100 mM EDTA) supplemented with 1% SDS, followed by incubation at 65°C for 30 min. Three hundred µL of 3 M potassium acetate was added to the mixture, which was subsequently placed on ice for 30 min. After centrifugation at 13,000 rpm for 20 min at 4°C, the supernatant (∼600 µL) was collected and mixed with a half volume of isopropanol. Samples were centrifuged at 13,000 rpm for 10 min, and the pellet was washed with 70% ethanol. Precipitates were dried and dissolved in 500 µL of TE buffer. Samples were then treated with RNase A (0.4-0.8 mg/mL) at 37°C for 15 min. For purification, each sample was mixed vigorously with the same volume of PCI (phenol:chloroform:isoamyl alcohol = 25:24:1, v/v) (Millipore Sigma cat#516726).

After centrifugation at 13,000 rpm for 5 minutes, the aqueous upper layer was collected and mixed vigorously with the same volume of chloroform, followed by another centrifugation at 13,000 rpm for 5 minutes. The upper layer (∼400 µL) was further subjected to ethanol precipitation. The final precipitates obtained were dried and dissolved in 100 µL of TE buffer. The typical yield of genomic DNA extracted from 200 flies was 0.2-0.5 mg. 10-20 ug of genomic DNA per genotype was digested with a restriction enzyme (BglII for characterizing the *TkR86C^LexA^* allele, XhoI for characterizing the *TkR99D^LexA^* allele) at 37°C overnight. Electrophoresis was performed using a 0.7% agarose gel. Roche Digoxigenin (DIG)-labeled DNA Molecular Weight Marker III (Millipore Sigma can# 11218603910) was loaded as a marker. The gel, placed on a shaker within an empty pipette tip box, was sequentially subjected to depurination (in 0.25 N HCl for 10 minutes), denaturation (in 0.5 M NaOH, 1.5 M NaCl for 15 minutes × 2), neutralization (in 0.5 M Tris/HCl (pH 7.5), 1.5 M NaCl for 15 minutes × 2), and equilibration (in 20X SSC for 10 minutes). DNA was transferred to a nylon membrane (Millipore Sigma cat# 11209299001) overnight by sandwiching the gel and membrane between paper towels soaked in 20X SSC under a 1.5 kilogram weight. DNA was immobilized onto the membrane by using a Stratalinker 2400 UV cross-linker.

DIG-labeled DNA probes were synthesized using Roche PCR DIG Probe Synthesis Kit (Millipore Sigma ca#11636090910). Probes were designed to target either the LexA coding sequence (Probe 1: 660-1,280 bp downstream from the start codon of the nls:LexA:p65) or the flanking genomic region specific for each gene. For *TkR86C*, the probe (Probe 2) was targeted to the genomic region 2,054 – 1,733 bp upstream of the 5’ end of the exon 1. For *TkR99D*, the probe (Probe 3) was targeted to the genomic region 1,814 – 2,317bp downstream from the 3’ end of the exon 2 (see full sequences of probes below). The DIG-labeled probes were hybridized to the membrane in Roche DIG Easy Hyb hybridization buffer (Millipore Sigma cat# 11603558001) at 49°C overnight. The membrane was sequentially washed twice with a low stringency buffer (2X SSC, 0.1% SDS) at room temperature for 5 minutes, and then twice with a pre-warmed high stringency buffer (5X SSC, 0.1% SDS) at 68°C for 15 minutes. After another brief wash with a DIG Easy Hyb kit wash buffer, the membrane was soaked in a DIG Easy Hyb blocking buffer at 4°C overnight. Roche anti-DIG-alkaline phosphatase Fab fragment (Millipore Sigma cat# 11093274910) were added to the blocking buffer at 1:10,000, and the membrane was incubated at room temperature for 30 min. The membrane was washed with the wash buffer for 15 min, twice, followed by a brief equilibration in a DIG Easy Hyb kit detection buffer. As a chemiluminescence substrate, Roche CDP-Star (Millipore Sigma cat#11759051001) was freshly diluted to 1:200 in the same buffer. Signals were developed on autoradiography films (Genesee Scientific #30-507).

### Animal preparation

Experimental flies for both behavioral and imaging experiments were collected on the day of eclosion into vials containing standard cornmeal-based food, and were kept as a group of up to 20 flies per vial at 25°C with 60% relative humidity and under a 9AM:9PM light:dark cycle. Flies used in *shibere^ts^* experiments were kept at 18°C. Tester flies were transferred to an aluminum foil-covered vial with food containing 0.2 mM all-*trans* retinal (Millipore Sigma, cat# R2500, 20 mM stock solution prepared in 95% ethanol) 5-6 days before experimentation. Every 3 days, flies were transferred to vials containing fresh food. Tester flies were aged for 5-7 days if carrying *Otd-nls:FLPo*, and 14-16 days if carrying *fru^FLP^*, to ensure consistent labeling of Tk-GAL4^FruM^ neurons (Asahina et al., 2014; Wohl et al., 2020). Rearing conditions of flies that carry *trans*-Tango elements are described below.

In behavioral experiments, a transgenic tester fly was paired with a “target” fly. Male target flies, wild type Canton-S individuals (originally from the lab of Martin Heisenberg, University of Würzberg), were group-reared with other male as virgins. To prepare mated female target flies, 5 Canton-S males were introduced into vials with ten, 4-days old virgin females and were reared for 2 more days to let them mate. The males used for mating were discarded. At 3-days old, both male and mated female target flies were briefly anesthetized with CO_2_, and the tip of one of their wings was clipped with a razor blade to distinguish them from tester flies when tracking. This clipping treatment did not reduce the amount of lunging detected under our experimental settings (data not shown).

### Behavioral assays

Behavior assays were conducted in the evening (from 4 to 9PM) at 22-25°C. For *shibere^ts^* experiments, flies were acclimated for 30 minutes at the temperatures of 22 or 32°C before testing. These experiments were performed in a climate-controlled booth kept at 60% relative humidity.

Social behavior assays were performed in a “12-well” acrylic chamber (Asahina et al., 2014) with food substrate (apple juice (Minute Maid) supplemented with 2.25% w/v agarose and 2.5% w/v sucrose; (Hoyer et al., 2008) covering the entire arena floor. The wall was coated with Insect-a-Slip (Bioquip Products, Inc., cat# 2871C) and the ceiling was coated with Surfasil Siliconizing Fluid (ThermoFisher Scientific, cat# TS-42800), to prevent flies from climbing as described previously (Asahina et al., 2014; Hoyer et al., 2008). Recording was done with USB3 digital cameras (Point Grey Flea3 USB3.0, FLIR Inc., cat# FL3-U3-13Y3M-C) controlled by the BIAS acquisition software (IORodeo, CA; https://bitbucket.org/iorodeo/bias). The camera was equipped with a machine vision lens (Fujinon, cat# HF35HA1B) and an infrared longpass filter (Midwest Optical Systems, cat# LP780-25.5) to block light from the LED sources used for optogenetic neuronal activation (see below). Movies were taken at 60 frames per second in the AVI format. Flies were discarded after each experiment. The food substrate was changed after 5 recordings.

To optogenetically activate Tk-GAL4^FruM^ neurons, a combined infrared (850nm) and optogenetic (625nm) LED backlight panel (described in https://www.janelia.org/open-science/combined-infrared-and-optogenetic-led-panel) was used as the light source. Briefly, the LED board was screwed to an aluminum heat sink (Aavid Thermalloy cat# 601403B06000) with a non-conductive thermal pad wedged between. Atop the board was a square wall of mirrors that faced inwards with 114 mm sides × 25 mm height. This mirror box was designed to ensure that light collected towards the edges of the board were similar in power to that collected towards the center of the board where more LEDs were present. Two 13mm thick acrylic plates, separated by 6mm, were placed above the backlight panel supported by 76mm optical poles. The first of the two plates was translucent white, which evenly diffused evenly the point source LEDs. An indicator infrared LED (850nm) was placed above the first plate to report optogenetic LED stimulation, which was invisible in the recorded videos due to longpass filter installed in front of the camera. The second plate was clear; fly behavior chambers rested upon it so that they were 25mm above the LED board. To minimize red light exposure before experiments, overhead fluorescent lights were covered in blue cellophane (Joann Fabrics cat# zprd_17968611a). Additionally, a black box surrounded the arena and LED backlight panel to keep out light from surrounding experiments. An opening on top of the box allowed optical access by camera as well as ambient light. It also had a small opening on one side to allow fly chambers to be moved in and out of the arena. The LED backlight panel was connected to a Teensy board, which interfaced with the flyBowl MATLAB custom code (kindly provided by Yoshi Aso and Jinyang Liu, HHMI Janelia Research Campus) so that the LEDs used for optogenetics were synchronized with the BIAS encoding software.

### Quantification of social behavior data

Acquired movies were analyzed largely as described in (Ishii et al., 2020; Leng et al., 2020; Wohl et al., 2020). In brief, the movies were first processed by the FlyTracker program (Eyjolfsdottir et al., 2014) (http://www.vision.caltech.edu/Tools/FlyTracker/). Number of lunges were quantified using behavioral classifiers developed in JAABA (Kabra et al., 2013) (https://source-forge.net/projects/jaaba/files/), as described in (Leng et al., 2020). The duration of time a tester fly orients toward a target fly (“time orienting”) was quantified as described previously (Ishii et al., 2020; Wohl et al., 2020). The distance traveled by a fly was calculated directly from the trx.mat file created by FlyTracker. The frame in which the infrared indicator LED turned on during the first LED stimulation period was used to align frames of movies.

### Immunohistochemistry

The following antibodies were used for immunohistochemistry with dilution ratios as indicated: rabbit anti-DsRed (1:1,000, Clontech cat# 632496, RRID: AB_10013483), mouse anti-BRP (1:100; Developmental Studies Hybridoma Bank nc82 (concentrated), RRID: AB_2314866), chicken anti-GFP (1:1,000, Abcam cat# ab13970, RRID:AB_300798), rat anti-HA (1:1000, Millipore Sigma cat#11867423001, RRID:AB_390918), goat anti-chicken Alexa 488 (1:100, ThermoFisher Scientific cat# A11039, RRID:AB_2534096), goat anti-rat Alexa 488 (1:100, ThermoFisher Scientific cat# A11006, RRID:AB_2534074), goat anti-rabbit Alexa 568 (1:100; ThermoFisher Scientific cat# A11036, RRID:AB_10563566), goat anti-mouse Alexa 633 (1:100; ThermoFisher Scientific cat# A21052, RRID:AB_2535719).

Immunohistochemistry of fly brains followed the protocol described in (Ishii et al., 2020; Wohl et al., 2020). Z-stack images were acquired by FV-1000 confocal microscopy (Olympus America) and were processed with Fiji software (Schindelin et al., 2012) (RRID:SCR_002285; https://fiji.sc/). Minimum and maximum intensity thresholds were adjusted for enhanced clarity. Registration of brains to the JCR2018 INTERSEX template brain (Bogovic et al., 2020) was performed as described in (Ishii et al., 2020; Jefferis et al., 2007; Wohl et al., 2020).

*Trans*-Tango flies used for immunohistochemistry were reared for 28-30 days at 21°C to allow sufficient expression of reporters in downstream areas with a maximal signal-to-noise ratio (Talay et al., 2017). To restrict expression of the human glucagon ligand, necessary for reporter translocation, to Tk-GAL4^FruM^ neurons, *Tk-GAL4^1^* expression was limited by a *tubulin-FRT-GAL80-FRT-stop* transgene and *fru^FLP^*.

### Image segmentation and quantification

To quantify the immunohistochemical fluorescence intensity of Syt:GFP and DenMark, Tk-GAL4^FruM^ neurons were first seqmented into the SMP projection, ring adjacent region, and axonal tract based on the confocal image of reporter proteins that visualize the neuroanatomy of Tk-GAL4^FruM^ neurons (myr:tdTomato for Syt:GFP samples, and cytosolic GFP for DenMark). The 3D-rendered images of Tk-GAL4^FruM^ neurons were manually segmented using the Paint Brush function of FluoRender software (Wan et al., 2009) as previously described in (Ishii et al., 2020; Wohl et al., 2020). Each segmented domain was converted back to an 8-bit stacked TIFF image, and a binary mask for the entire stack was created by adjusting the threshold value (20-40 depending on the image quality) in ImageJ. The average signal intensity within the given domain was calculated as [sum of signal intensity in pixels within the mask]/[total number of pixels within the mask].

Signal intensity of GCaMP6f immunohistochemical fluorescence of *TkR86C^LexA^* and *TkR99D^LexA^* neurons in the vicinity of Tk-GAL4^FruM^ neurons was calculated in a similar manner as above. The SMP projection and ring adjacent region were segmented based on the confocal image of CsChrimson:tdTomato expressed in Tk-GAL4^FruM^ neurons.

### Functional Imaging

On the day of the experiment, flies were briefly anesthetized on ice and mounted on a custom chamber using ultraviolet curing adhesive (Norland Optical Adhesive 63, Norland Products, Inc.) to secure the head and thorax to a tin foil base. The proboscis was also dabbed with glue to prevent its extension from altering the position of the brain. The head cuticle was removed with sharp forceps in *Drosophila* adult hemolymph-like saline (Wang et al., 2003) at room temperature. After cuticle removal, the saline was exchanged with a fresh volume.

Optogenetic stimulation was applied with an external fiber-coupled LED of 625 nm (Thorlabs, cat# M625F2) controlled by a programmable LED driver (ThorLabs, cat# DC2200). The end of the LED fiber (Thorlabs, cat# M28L01) was placed 5 mm from the brain. The LED produced 10-millisecond pulses 10 seconds at 0.5, 1, or 5 Hz. The energy from the LED that the neurons received was estimated from the measurement of the LED power as 0.2 mA using a photodiode power sensor (Thorlabs, cat# S130C) coupled to a digital optical power/energy meter (Thorlabs, cat# PM100D) 5 mm away from the end of the LED fiber.

The multiphoton laser scanning microscope (FV-MPE-RS, Olympus Corporation), equipped with 25X water immersion objective (Olympus Corporation, cat# XLPLN25XWMP2), was used for monitoring the fluorescence of GCaMP6f. The recordings began 5-10 seconds before a 10 second stimulation and continued for 10-20 seconds after stimulation for a total of 25-40 seconds. GCaMP6f fluorescence was visualized with a tunable laser set at 920 nm output (Spectra-Physics Insight DL Dual-OL, Newport Corporation), and CsChrimson:tdTomato was visualized with an auxiliary laser with a fixed output of 1040 nm. Images were taken at 5-7 Hz, depending on the size of scanning area, with a 256 × 256 pixel resolution.

Acquired images (.OIR format) were converted and analyzed in Fiji with the Olympus ImageJ plugin (http://imagej.net/OlympusImageJPlugin). Imaging windows were chosen that maximally captured the Tk-GAL4^FruM^ neuronal projections in the SMP or in the ring adjacent region using the fluorescence of CsChrimson:tdTomato. Polygonal regions of interest were drawn using the tdTomato fluorescence, and ΔF/F of GCaMP6f was calculated using a custom-written MATLAB code. First, the baseline fluorescence value (F_base_) was calculated by averaging the fluorescence for 5 seconds preceding the stimulation. ΔF/F for each frame ((ΔF/F)_frame=N_) was calculated as follows:

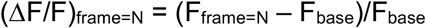

Then, the ΔF/F_frame=N_ for frames taken during the 10-second LED stimulation were averaged to calculate the ΔF/F of a given trial. Frames that contained LED light for optogenetic stimulation were excluded from the analysis. Values from 1-5 trials were averaged for each condition. Trials with excessive movement were discarded.

Our preliminary study indicated that baseline fluorescence of the *QUAS-GCaMP3* transgene (Bloomington #52231) driven by *trans*-Tango was not sufficient to be visualized under 2-photon microscopy. Thus, we constructed *15XQUAS-IVS-Syn21-GCaMP6f-p10* (see above for details) and used two copies of the insertions for *trans*-Tango imaging experiments. These flies were transferred to 0.2 mM all-*trans*-retinal food six days before experimentation. Due to the higher level of expression of our GCaMP6f constructs, we needed to age flies only for 16-20 days.

### Pharmacology

A 2.5 mM master solution of mecamylamine was made by dissolving mecamylamine hydrochloride (Millipore Sigma cat# M9020) in *Drosophia* adult hemolymph saline (Wang et al., 2003). Pre-treatment trials were recorded first. Then, mecamylamine saline was added to the imaging saline reservoir for a final concentration of 25 µM via pipetting. The drug-infused saline was then gently mixed. For vehicle experiments, the same amount of saline was added, but without mecamylamine. Imaging resumed 15 minutes after adding the solution. When “treatment” trials were complete, washout of drug was performed in the following steps. First, the saline, whether with or without mecamylamine, was replaced with drug-free saline 6 times. 15 minutes later, the saline was again replaced twice. Calcium imaging for the washout condition resumed 15 minutes after the second wash cycle so that it began a total of about 30 minutes after the first washing.

### Statistical Analysis

Statistical analyses were carried out using MATLAB except calculation of 95% confidence intervals, which was done using Microsoft Excel’s ‘CONFIDENCE.T’ function. Non-parametric analyses were used for behavioral data. After behaviors within each time window were calculated, the Kruskal-Wallis one-way ANOVA (‘kruskalwallis’) was used to evaluate whether a given behavior was significantly different among more than 2 different illumination periods or genotypes. When the p-value was below 0.05, the post-hoc Mann-Whitney signed rank test (‘signrank’) was used to detect significant differences between illumination period, and the Mann-Whitney U-test (‘ranksum’) was used to detect significant differences between genotypes. In both cases, Bonferroni correction was applied to p values. When the uncorrected p value was less than 0.05 but the corrected value did not pass the significant level, the uncorrected value was shown on panels with parentheses.

Fluorescence data from immunohistochemical and functional imaging data were analyzed using parametric tests. Datasets from more than 2 independent sources (e.g., different genotypes) were first analyzed with one-way ANOVA (‘anova1’). When the p-value was below 0.05, the post-hoc Welch’s t-test (‘ttest2’) with Bonferroni correction was used to detect significant differences between genotypes or conditions. Datasets from more than 2 balanced sources (e.g., multiple measurements from the same sample) were first analyzed with two-way ANOVA (‘anova2’). When the p-value was below 0.05, the post-hoc paired t-test (‘ttest’) with Bonferroni correction was used to detect significant differences between measurements. In both cases, ANOVA was omitted when comparing 2 datasets.

All data points, statistical results, and (for parametric tests) 95% confidence intervals are presented in Source Data Table.

## SUPPLEMENTAL FIGURES

**Figure S1:**
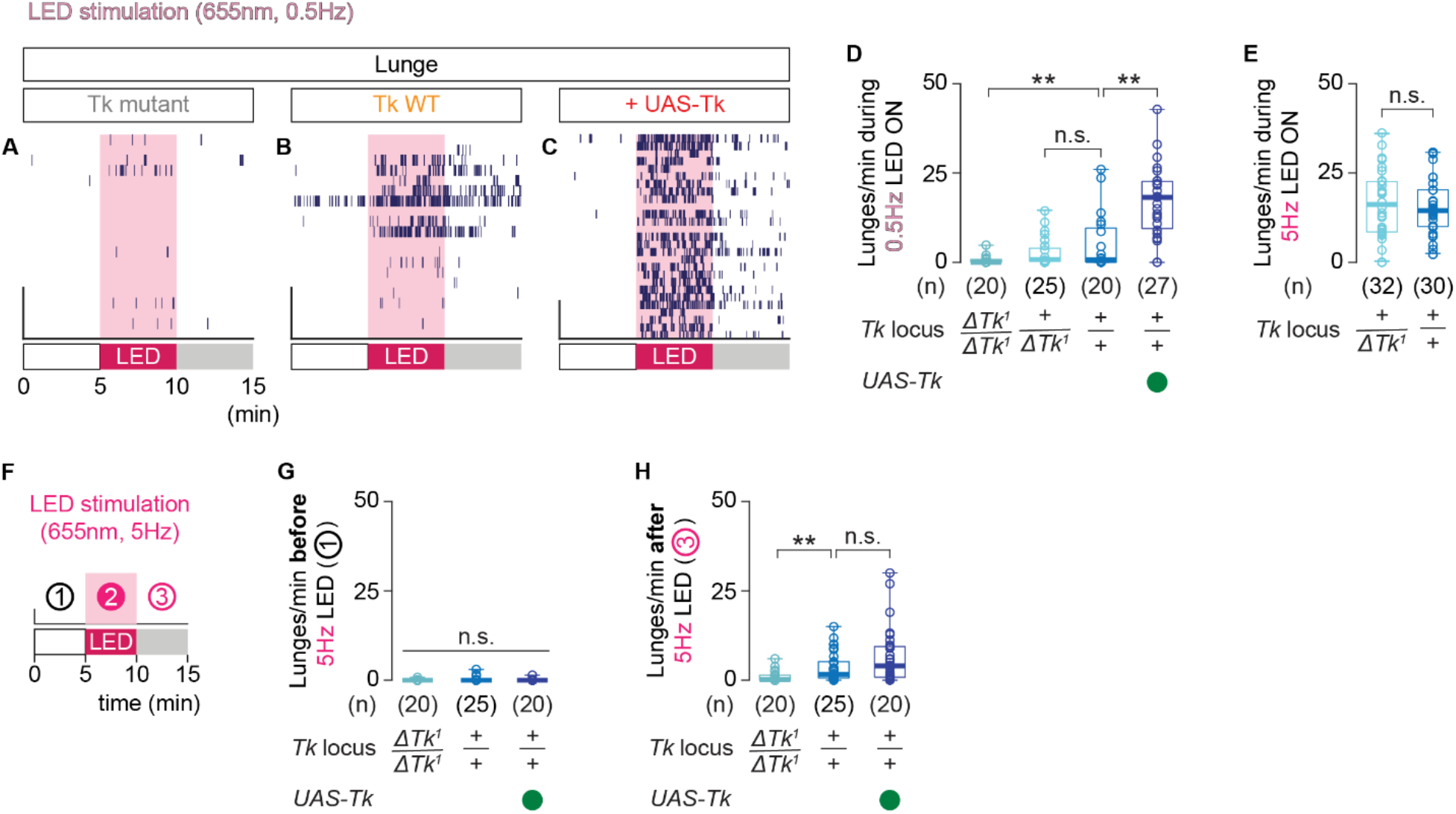
Tachykinin amount controls the intensity of aggression induced by optogenetic activation of Tk-GAL4^FruM^ neurons at low stimulation frequency. **A-C**. Raster plots of lunges induced by optogenetic activation of Tk-GAL4^FruM^ neurons at 0.5 Hz. **D**, **E**. Boxplots of lunges performed during optogenetic stimulation (**D**; 0.5 Hz, **E**: 5 Hz) of Tk-GAL4^FruM^ neurons. Genotypes of the *Tk* locus are indicated below. The *Tk* wild-type dataset in **E** is replotted from Fig. 1E. **F**. Schematic representations of behavioral assays with optogenetic stimulation. **G**, **H**. Boxplots of lunges performed before (**G**) or after (**H**) optogenetic stimulation of Tk-GAL4^FruM^ neurons (also see **F**). Target flies were group-housed wild-type males. In **D** and **H**, ** p < 0.01, n.s. > 0.05 by Kruskal-Wallis one-way ANOVA (**D**: p < 3.5 × 10^−9^, **H**: p < 0.002) and post-hoc Mann-Whitney U-test. In **E**, n.s. p > 0.05 by Mann-Whitney U-test. In **G**, n.s. p > 0.05 by Kruskal-Wallis one-way ANOVA.

**Figure S2:**
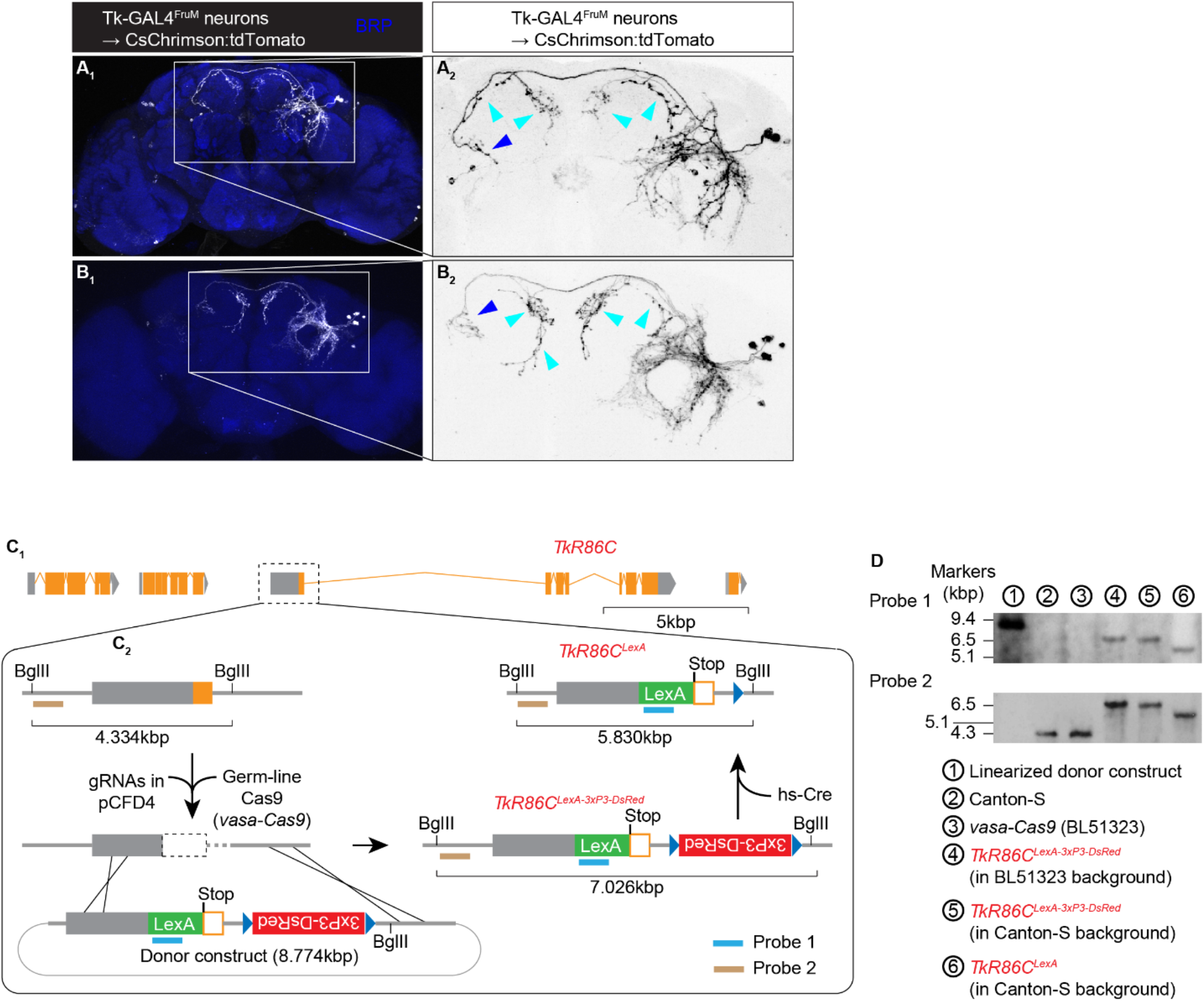
Details of Tk-GAL4^FruM^ neuroanatomy and the *TkR86C^LexA^* allele. **A**, **B**. Two independent samples of unilaterally labeled Tk-GAL4^FruM^ neurons are shown. These were generated by stochastic inactivity of fruFLP on one side of the brain in animals that also carried *Tk-GAL4^1^* and *20XUAS>myr:TopHAT2>CsChrimson:tdTomato*(in attP2) transgenes. Magnified images of Tk-GAL4^FruM^ neurons in **A_1_** and **B_1_** are shown in **A_2_** and **B_2_**, respectively. Cyan arrowheads indicate bouton-like varicosities in the SMP that enamate from the putative axon tract, which crosses the midline and reaches the area where the Tk-GAL4^FruM^ neurons in the contralateral side extend their dendritic arbors (blue arrowheads). **C**: A schematic of the steps taken to create the *TkR86C^LexA^* allele. The first exon of *TkR86C* was modified (**C_1_**) with CRISPR/Cas9-mediated homologous recombination as described in **C_2_**. **D**: Southern blotting analysis of *TkR86C^LexA^* alleles. Probe 1 targets upstream of the *TkR86C* locus whereas probe 2 targets the coding region of LexA (see **C_2_**). Note that flies 4–6 were homozygous for the *TkR86C* alleles.

**Figure S3:**
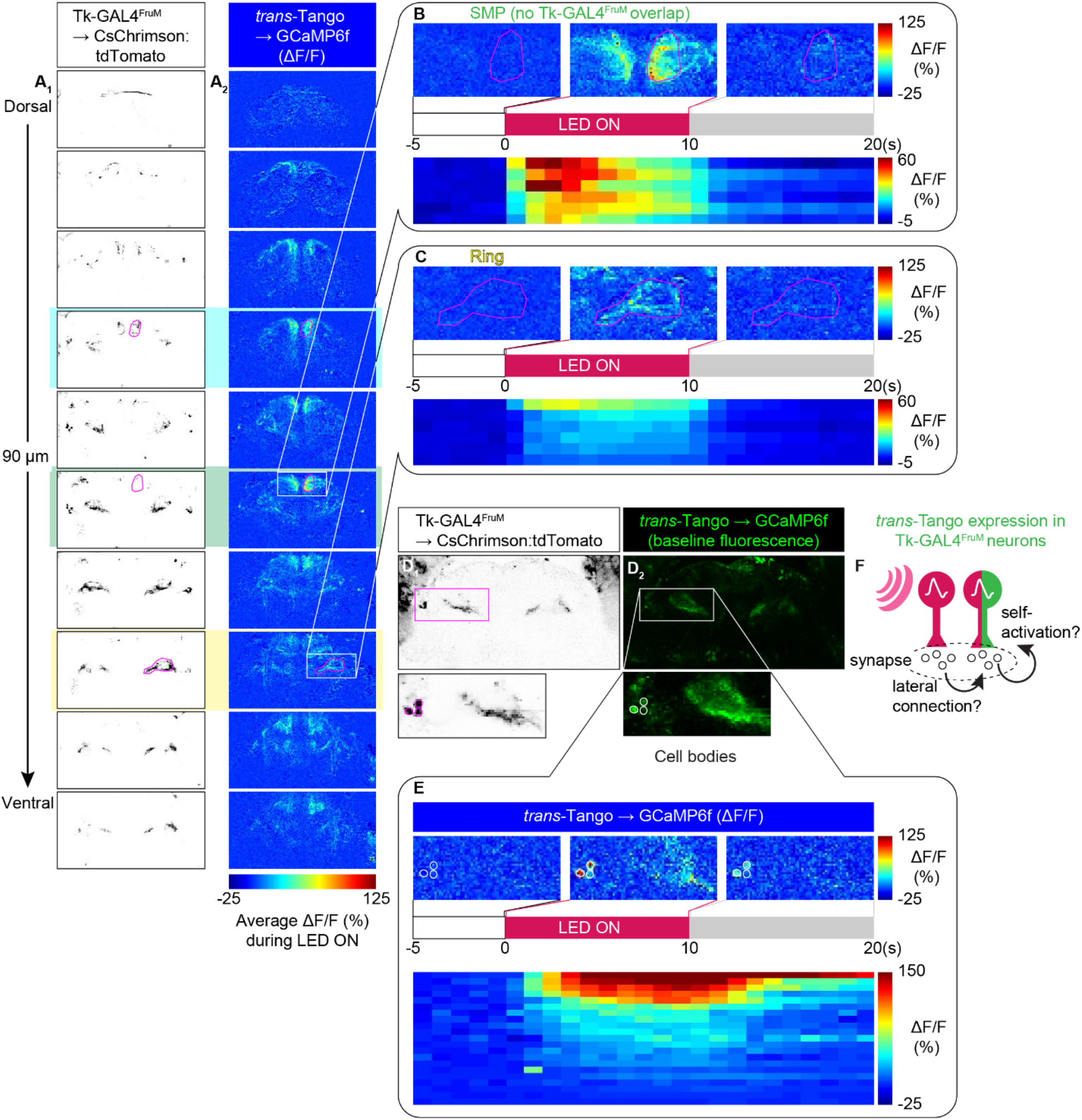
Details of neural activity from *trans*-Tango-labeled cells. **A.** Images of CsChrimson:tdTomato fluorescence in Tk-GAL4^FruM^ neurons (**A_1_**) and average normalized change in GCaMP6f fluorescence (ΔF/F) in putative synaptic downstream partners of Tk-GAL4^FruM^ neurons labeled by *trans*-Tango (**A_2_**) in a representative sample during optogenetic stimulation of Tk-GAL4^FruM^ neurons. **B, C**. Pseudocolored GCaMP6f fluorescence changes (ΔF/F) from the area posterior to the projection of Tk-GAL4^FruM^ neurons to the SMP (**B**) or the ring-adjacent region (**C**), represented as a spatio-temporal average from the sample shown in **A** (top) and as a time course binned into seconds from 6 independent samples (bottom). Unlike neurons labeled by *TkR86C^LexA^* (see Figure 3), calcium increased in the area near the projection of Tk-GAL4^FruM^ neurons to the ring-adjacent region. **D**. Images of CsChrimson:tdTomato fluorescence in Tk-GAL4^FruM^ neurons (**D_1_**) and baseline fluorescence of GCaMP6f in *trans*-Tango-labeled neurons (**D_2_**) in a representative sample. Some, if not all, Tk-GAL4^FruM^ neurons expressed GCaMP6f. **E**. Pseudocolored GCaMP6f fluorescence changes (ΔF/F) in cell bodies of Tk-GAL4^FruM^ neurons (**C**), represented as a spatio-temporal average from the sample shown in **A** (top) and as a time course, binned into seconds, from 21 cell bodies in 7 independent brains (bottom). **F**. *trans*-Tango may label Tk-GAL4^FruM^ neurons either via lateral connections among the group of cells, or via self-labeling due to the presence of the receptor component of *trans*-Tango in these cells. Note that the modified human glucagon receptor is expressed by the pan-neuronal promoter of N-synaptobrevin.

**Figure S4:**
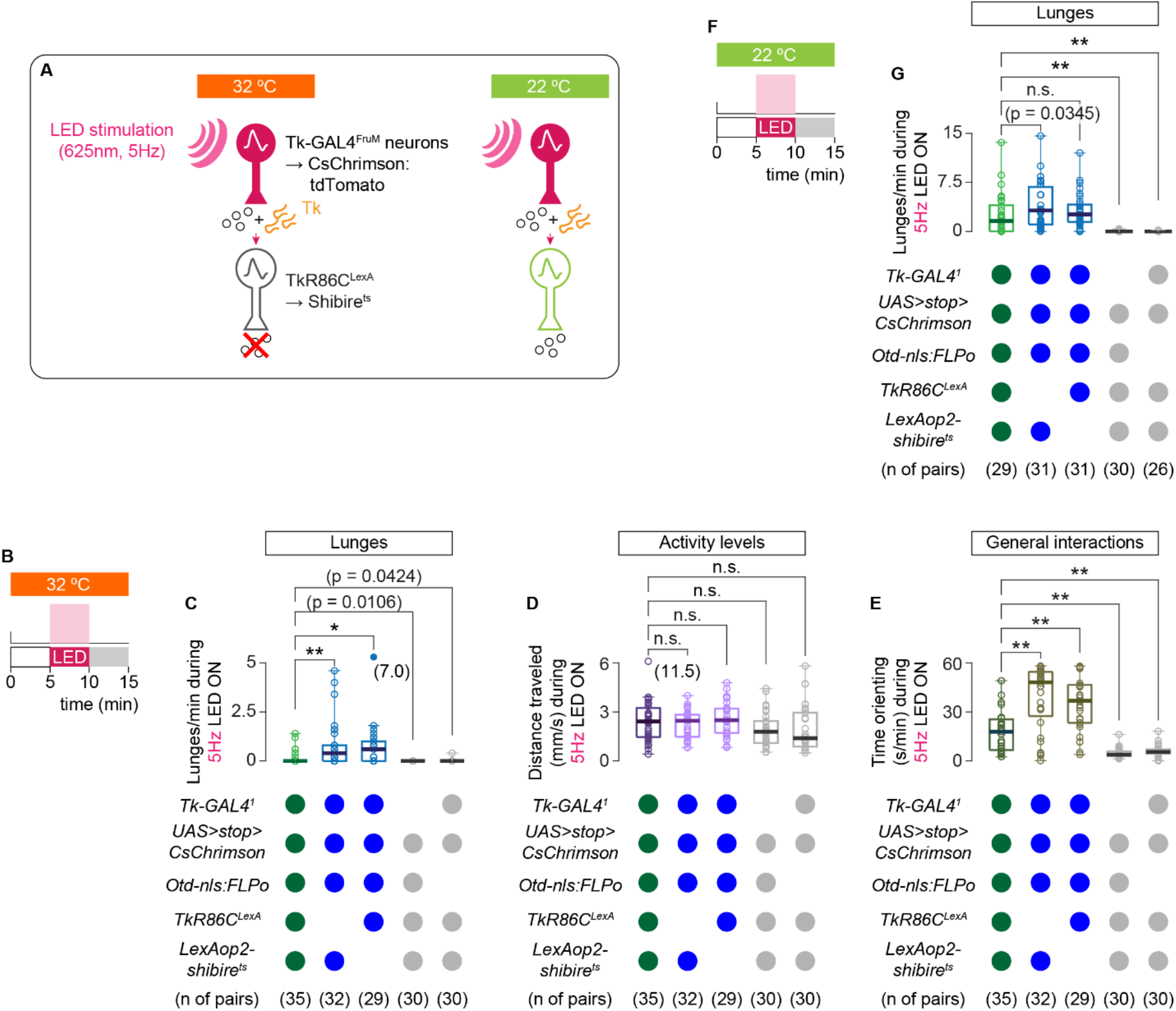
Blocking of synaptic transmission from *TkR86C^LexA^* neurons suppresses aggression induced by optogenetic activation of Tk-GAL4^FruM^ neurons. **A.** Schematic of the genetic manipulations used in the behavioral experiment in this figure. **B**. Design of the optogenetic behavioral experiments in the restrictive temperature of 32°C. **C.** Boxplot of number of lunges performed by the flies with the indicated genotypes during each time window. ** p < 0.01, * p < 0.05 by Kruskal-Wallis one-way ANOVA (p < 8.1 × 10^−13^) and post-hoc Mann-Whitney U-test. **D.** Boxplots of distance traveled by the same pairs of flies as in **C**. n.s. p > 0.05 by Kruskal-Wallis one-way ANOVA (p = 0.047) and post-hoc Mann-Whitney U-test. **E.** Boxplots of duration of orienting towards an opponent (the same pairs of flies as in **C**). ** p < 0.01 by Kruskal-Wallis one-way ANOVA (p < 2.4 × 10^−16^) and post-hoc Mann-Whitney U-test. **F**. Design of the optogenetic behavioral experiments in the permissive temperature of 22°C. **G.** Boxplot of number of lunges performed by the flies with the indicated genotypes during each time window. ** p < 0.01 by Kruskal-Wallis one-way ANOVA (p < 4.4X10^−18^) and post-hoc Mann-Whitney U-test.

**Figure S5:**
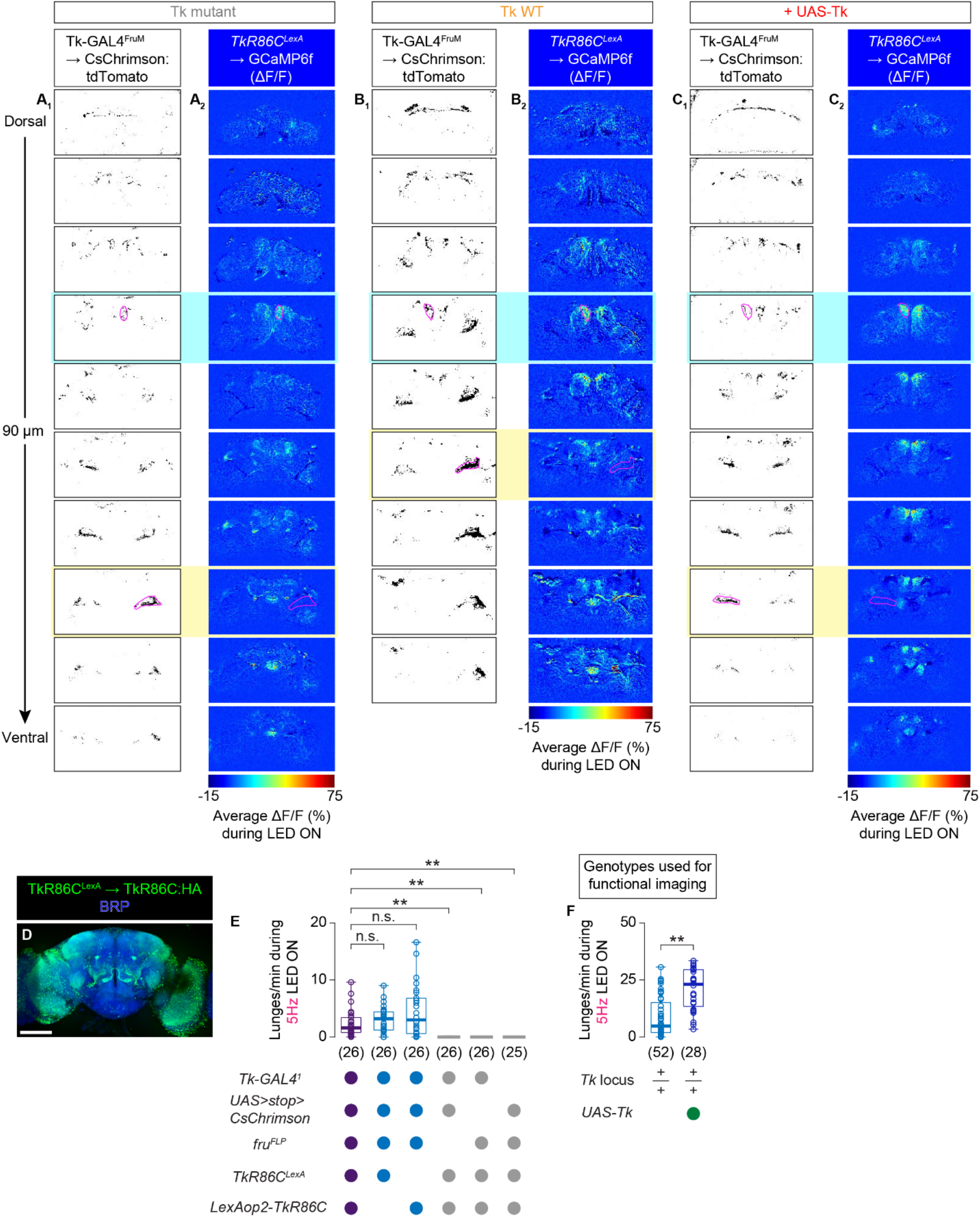
Manipulation of tachykinin and TkR86C on the function of Tk-GAL4^FruM^ neurons. **A-C.** Images of CsChrimson:tdTomato fluorescence in Tk-GAL4^FruM^ neurons (**A_1_-C_1_**) and average fluorescence of GCaMP6f in *TkR86C^LexA^* neurons (**A_2_-C_2_**) in a representative sample from *Tk* null mutants (**A**), *Tk* wild type (**B**), and in an animal with a *UAS-Tk* transgene (**C**). The GCaMP6f signals are shown in pseudocolor as the relative increase of fluorescence (ΔF/F) during optogenetic activation of Tk-GAL4^FruM^ neurons, as indicated below. **D**. Representative expression pattern of TkR86C tagged with HA (green) driven by *TkR86C^LexA^* in the brain along with the neuropil marker Brp, visualized by immunohistochemistry. Scale bar: 100 µm. **E**. Boxplots of lunges performed during optogenetic stimulation of Tk-GAL4^FruM^ neurons while TkR86C is over-expressed by *TkR86C^LexA^* (purple, right), along with genetic controls as indicated below. ** p < 0.01, n.s. p > 0.5 by Kruskal-Wallis one-way ANOVA (p < 4.6 × 10^−23^) and post-hoc Mann-Whitney U-test. **F**. Boxplots of lunges performed during optogenetic stimulation of Tk-GAL4^FruM^ neurons, with (right) or without (left) the *UAS-Tk* transgene, by the genotypes used for the functional imaging. ** p < 0.01 by Mann-Whitney U-test.

**Figure S6:**
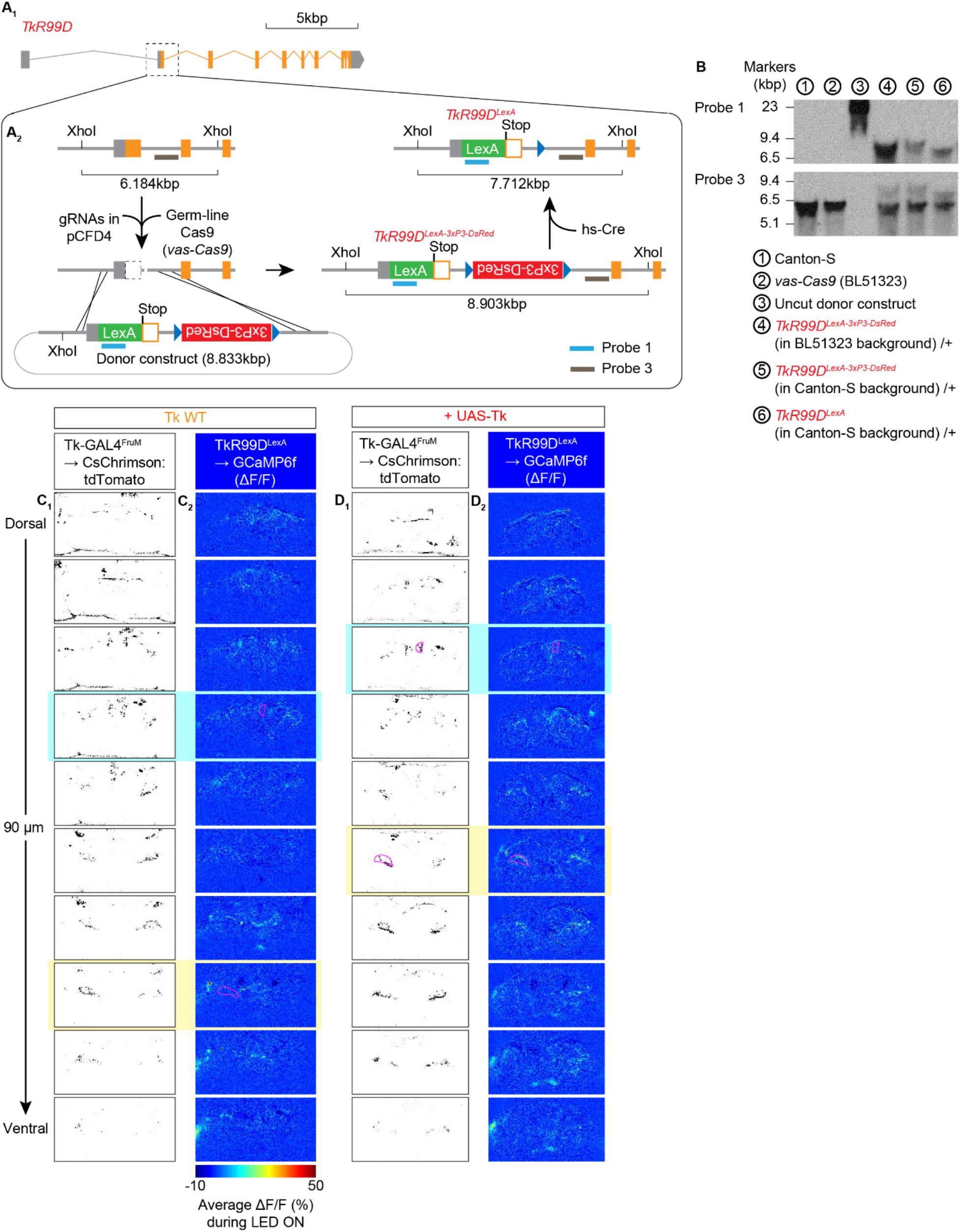
Characterization of the *TkR99D^LexA^* allele and neurons visualized by the allele. **A**. A schematic of the steps taken to create the *TkR99D^LexA^* allele. The second exon of *TkR99D* was modified (**A_1_**) with CRISPR/Cas9-mediated homologous recombination as described in **A_2_**. **B**. Southern blotting analysis of *TkR99D^LexA^* alleles. Probe 1 targets the coding region of LexA (also used in Fig. S1A-B) whereas probe 3 targets the area downstream of the *TkR99D* 2^nd^ exon (see **A_2_**). Note that flies 4-6 were heterozygous for the *TkR99D* alleles, therefore ∼6.2 kb fragments from the wild type allele were also present. **C**, **D.** Images of CsChrimson:tdTomato fluorescence in Tk-GAL4^FruM^ neurons (**C_1_-D_1_**) and average fluorescence of GCaMP6f in *TkR99D^LexA^* neurons (**C_2_-D_2_**) in a representative sample from a *Tk* wild type animal (**C**), and in an animal with a *UAS-Tk* transgene (**D**). The GCaMP6f signals are shown in pseudocolor as the relative increase in fluorescence (ΔF/F) during optogenetic activation of Tk-GAL4^FruM^ neurons, as indicated below. Frames with cyan and yellow backgrounds were used for measurements of the responses in the SMP and the ring-adjacent region, respectively.

## SUPPLEMENTAL TABLES

**Table S1:**
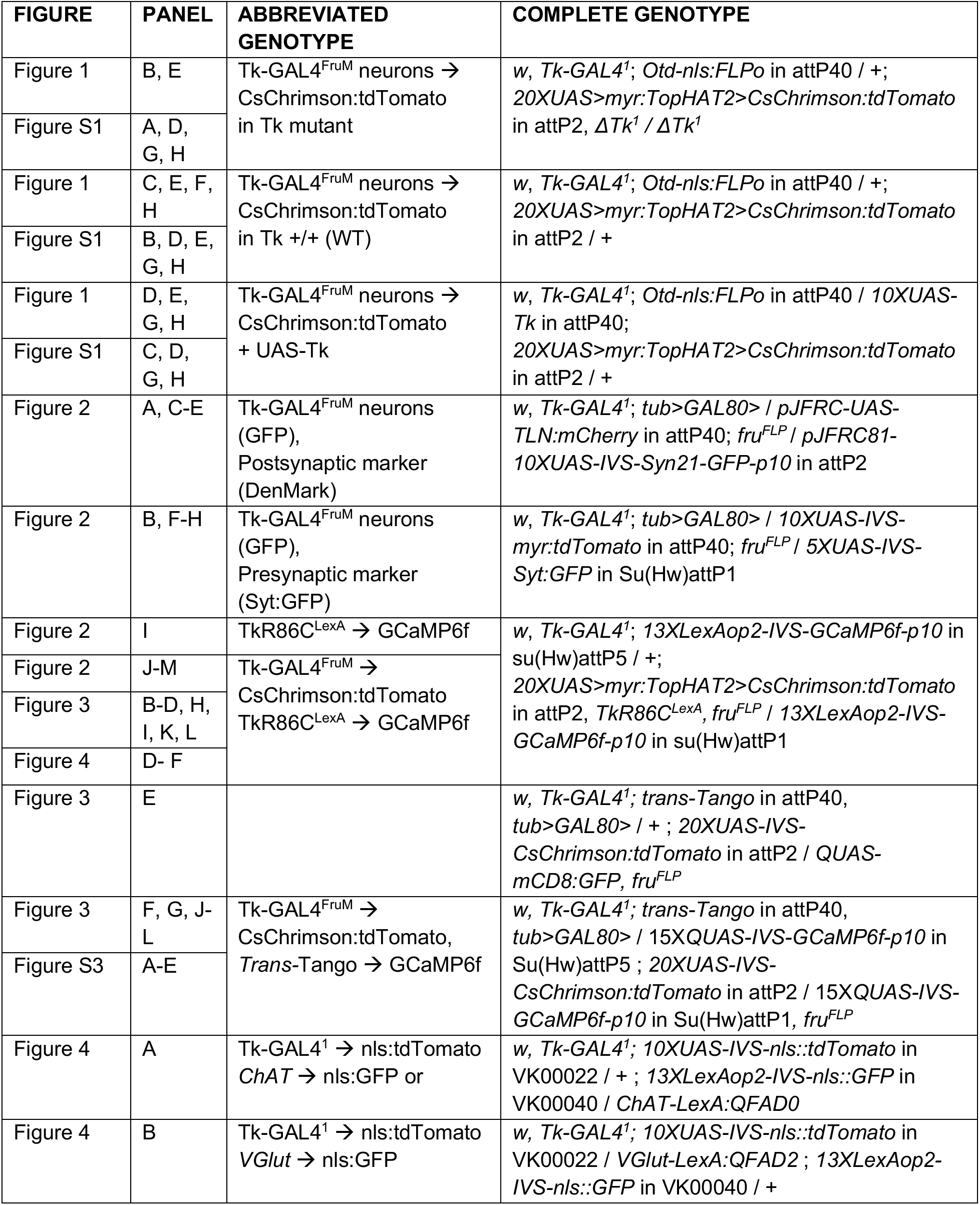
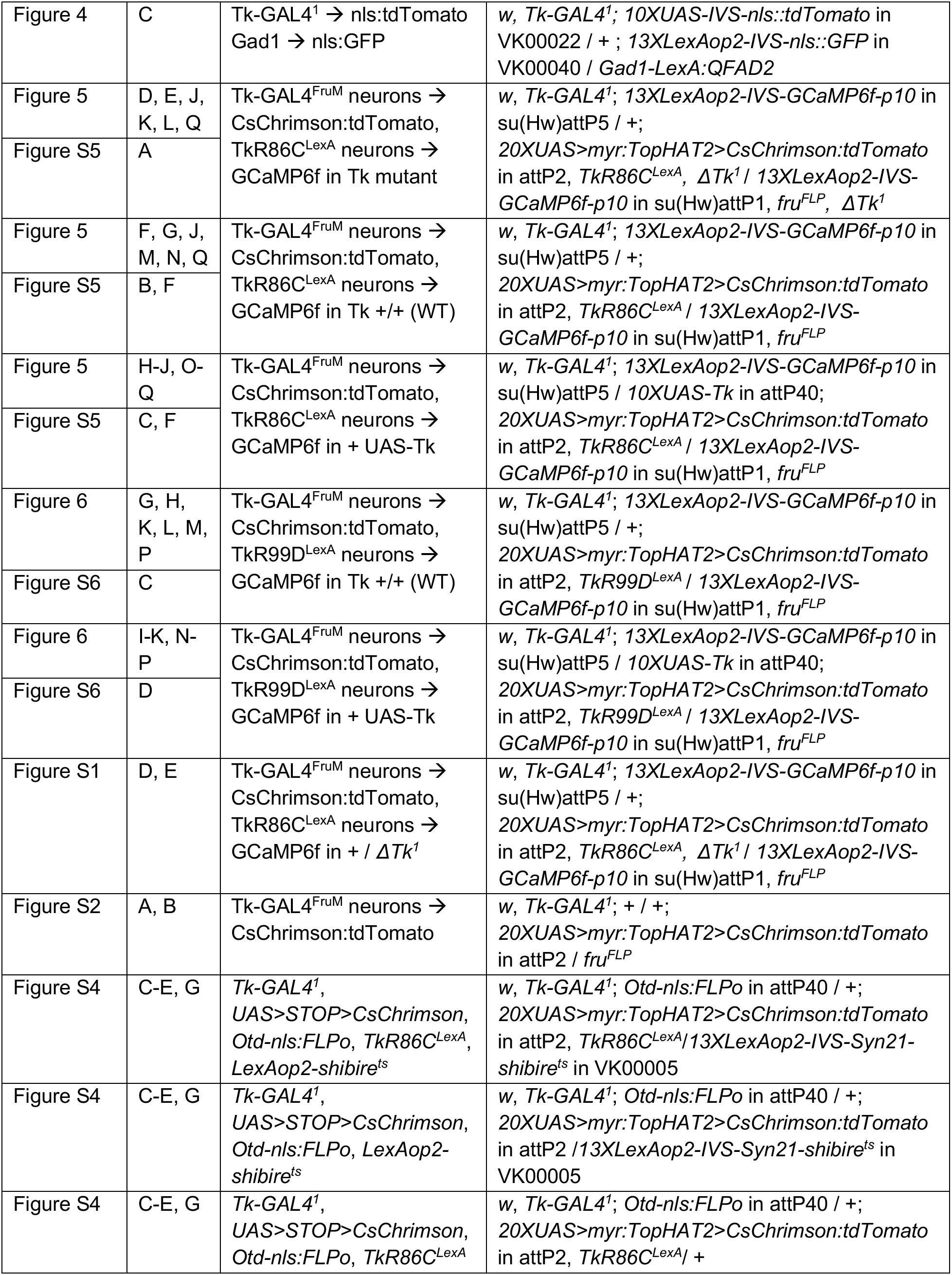
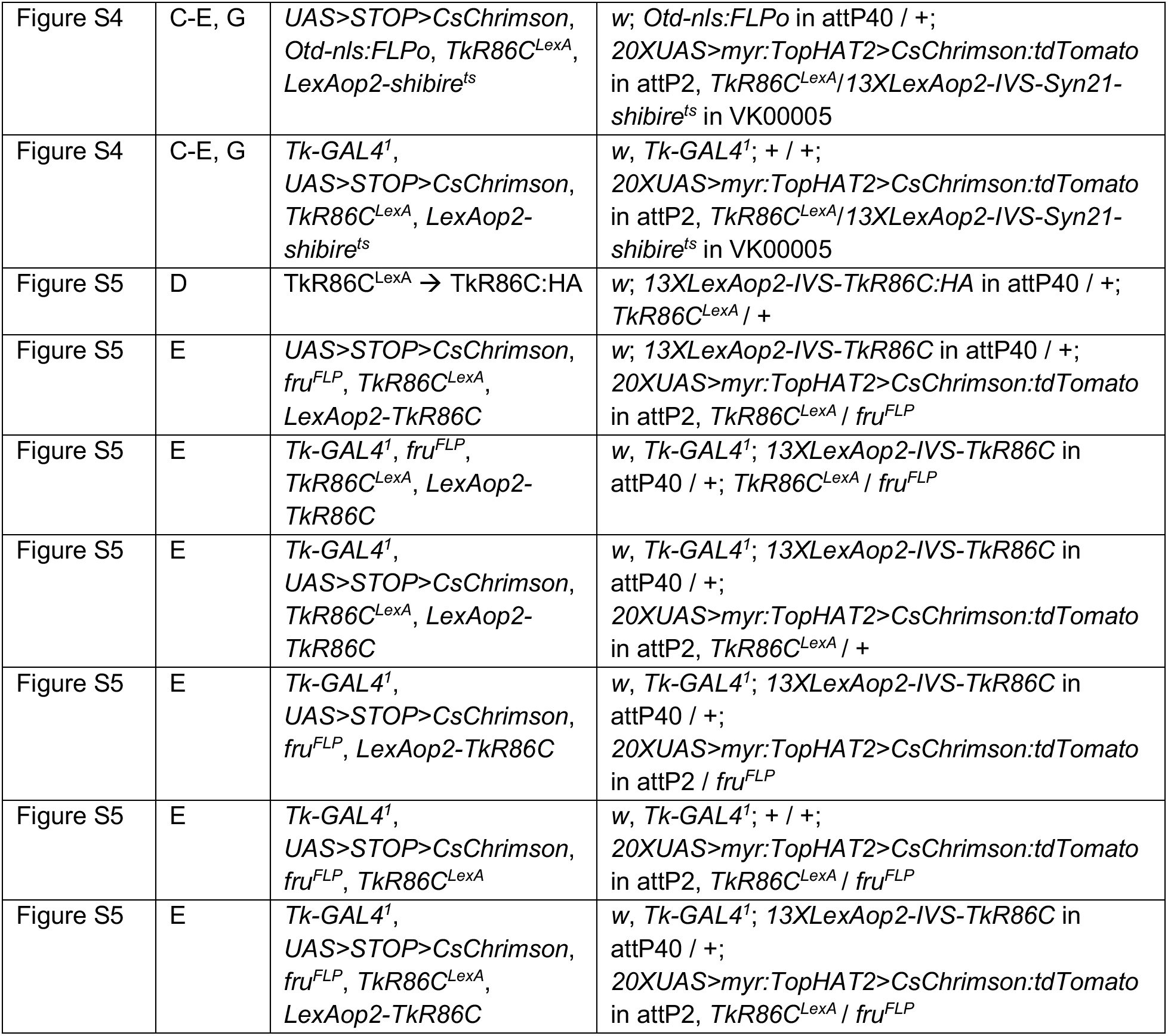
Complete genotypes of Drosophila strains used in this study, related to all Figures.

**Table S2:**
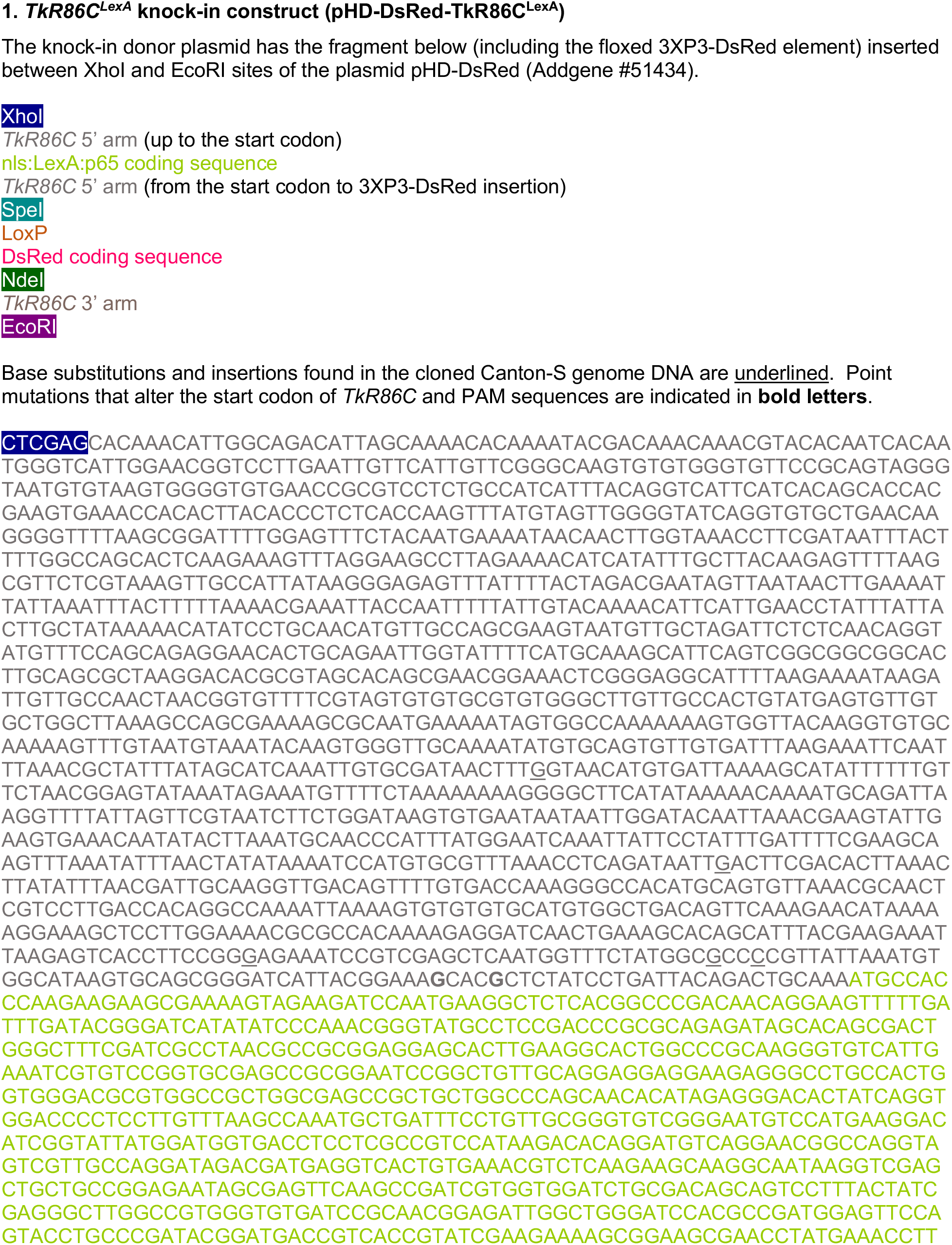
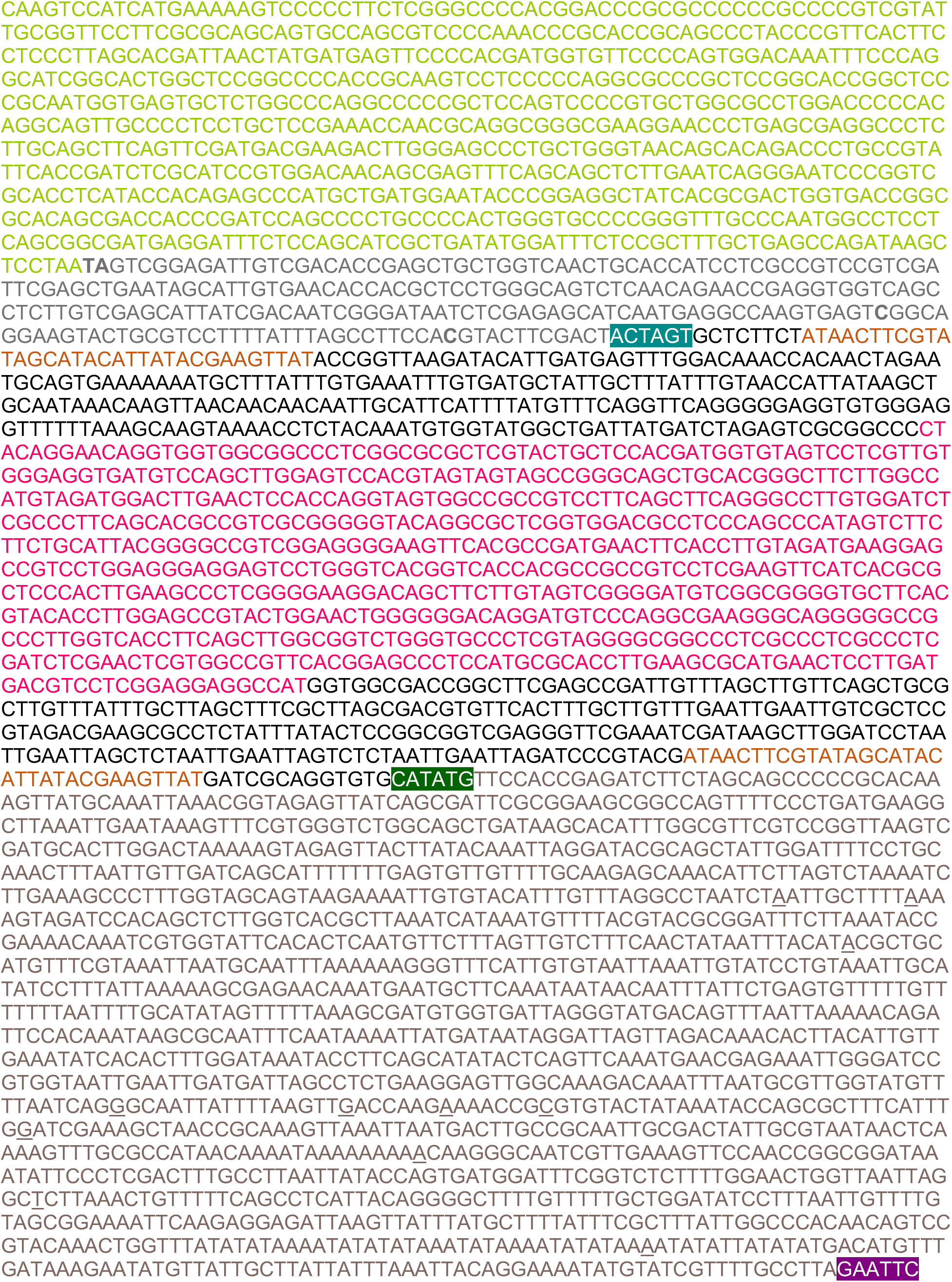
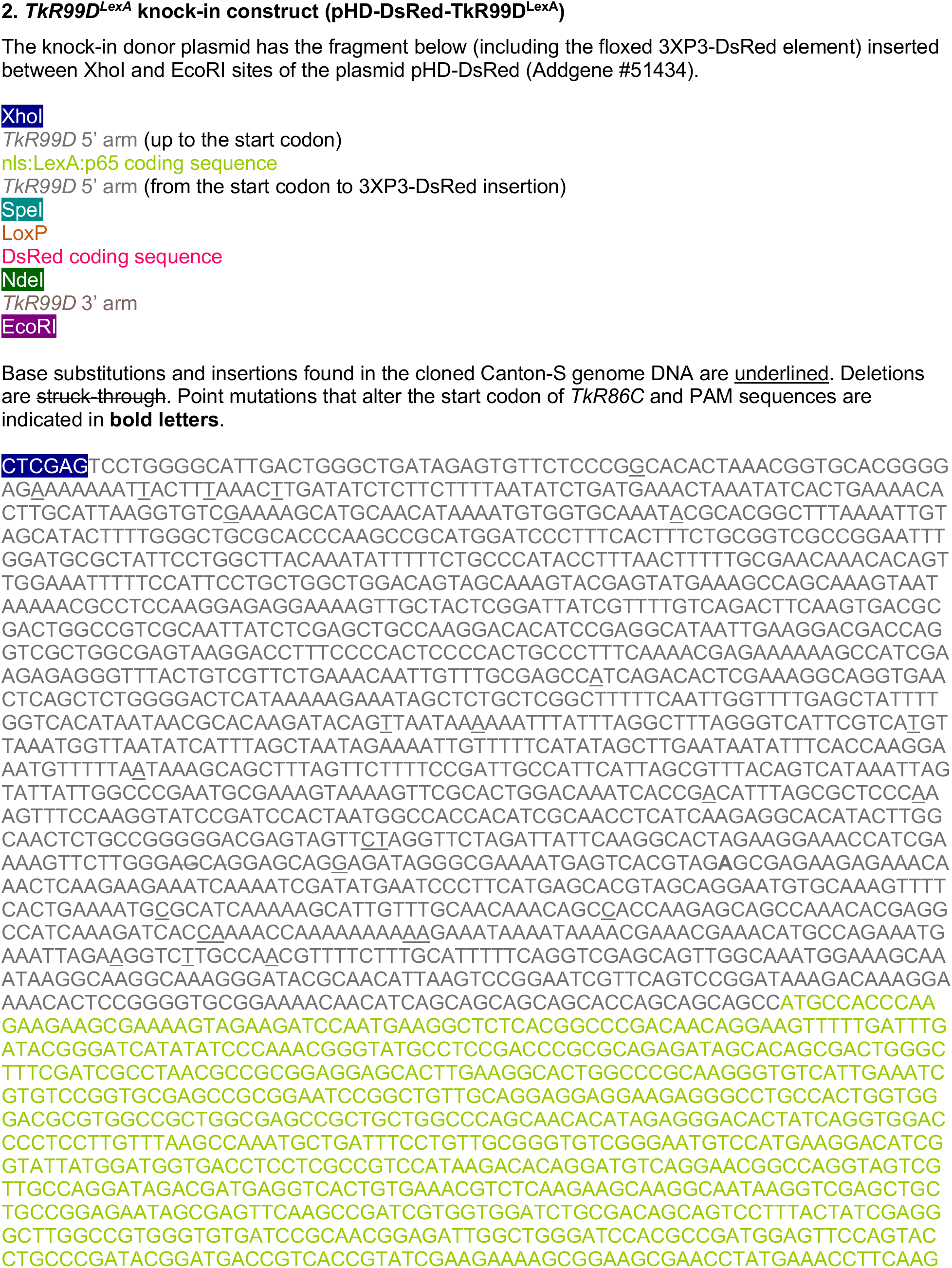
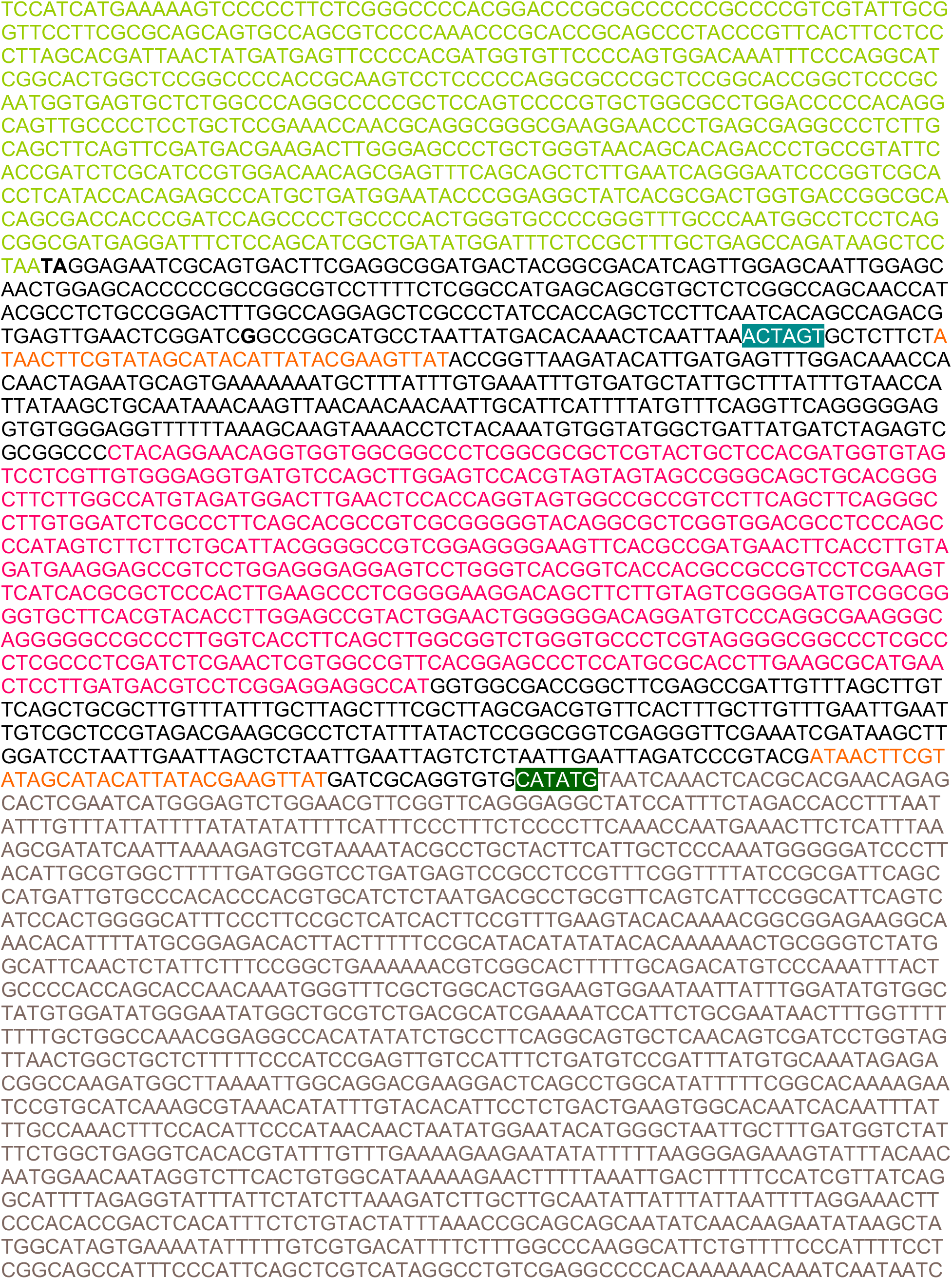
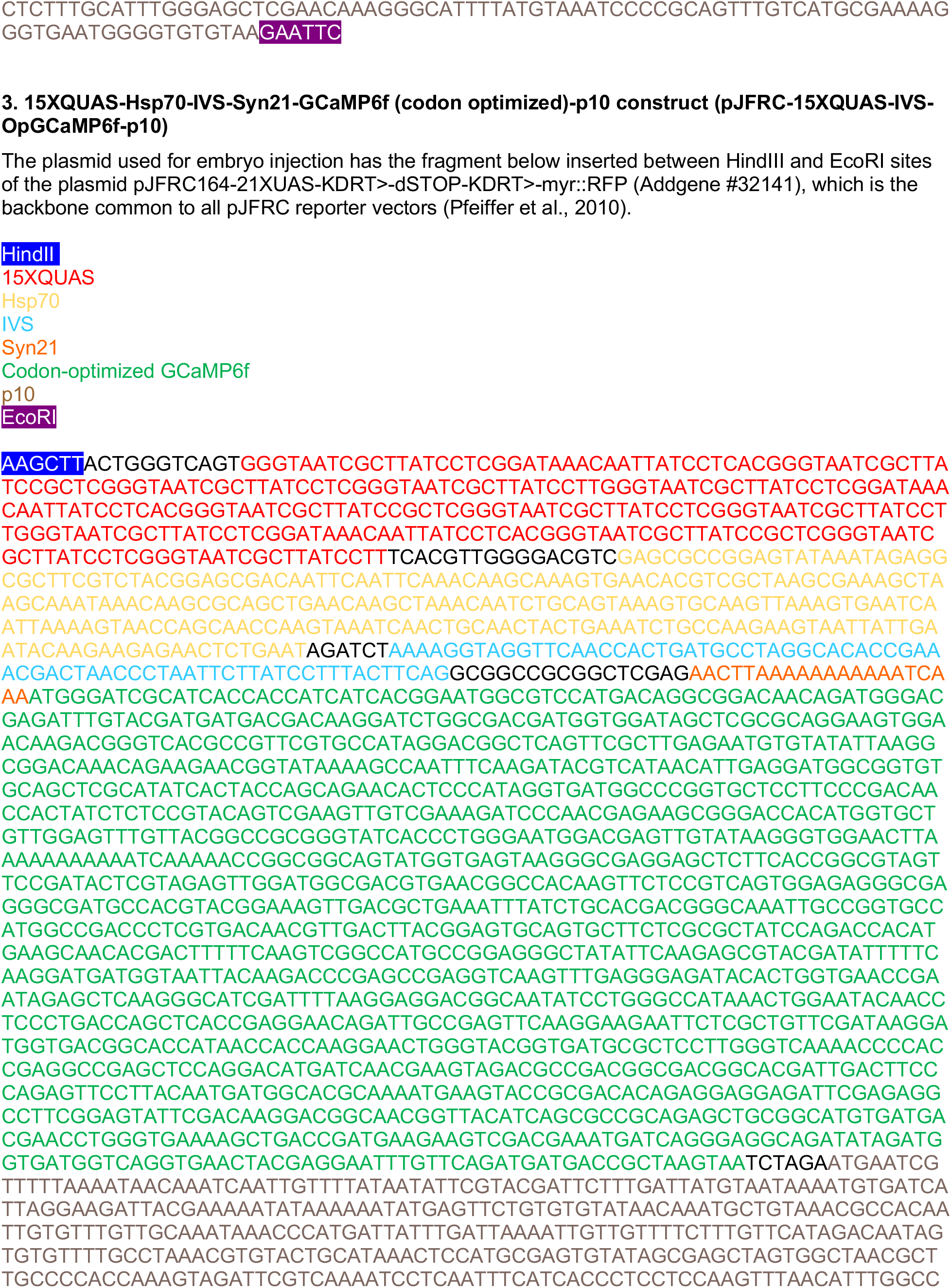
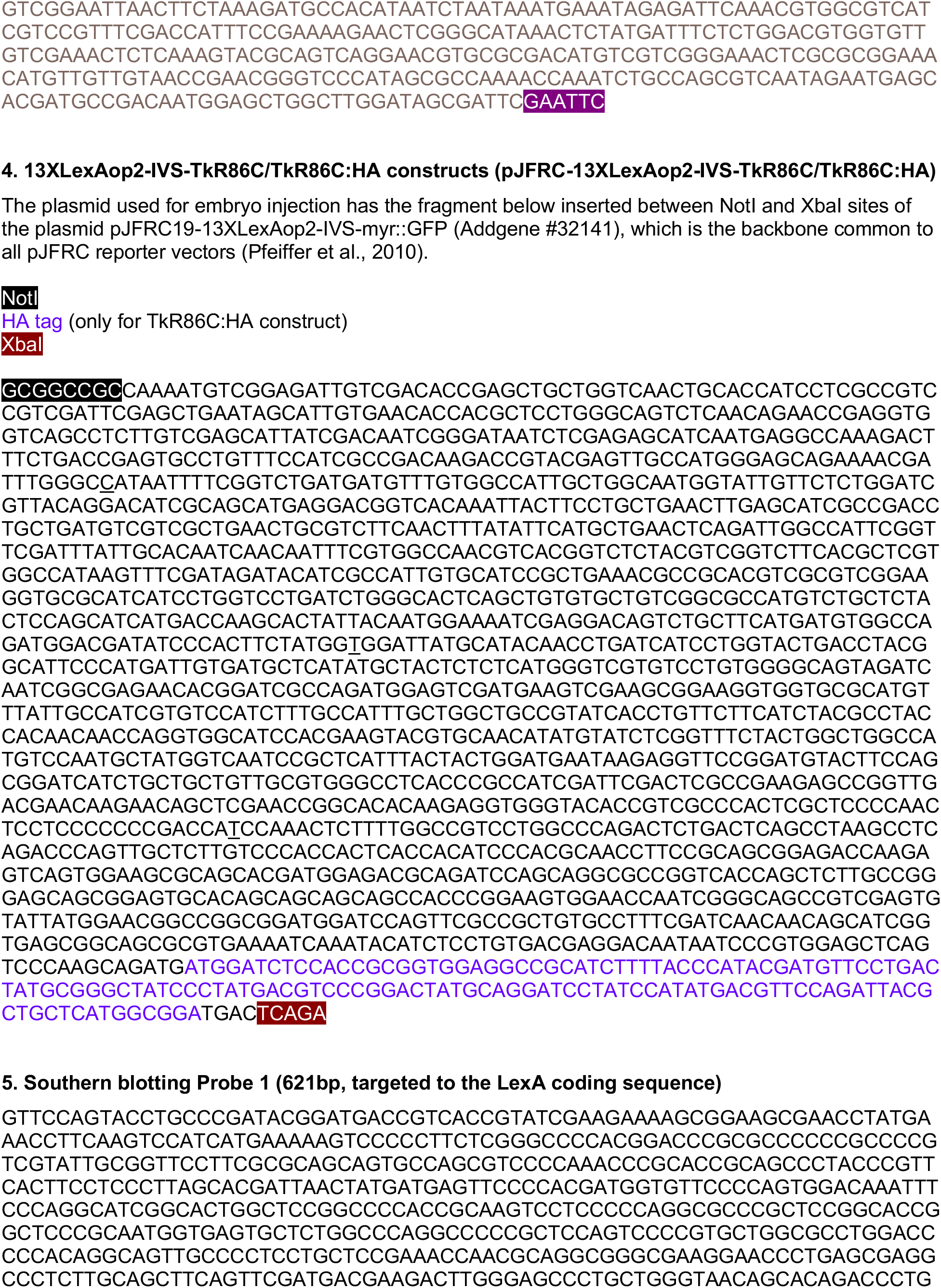
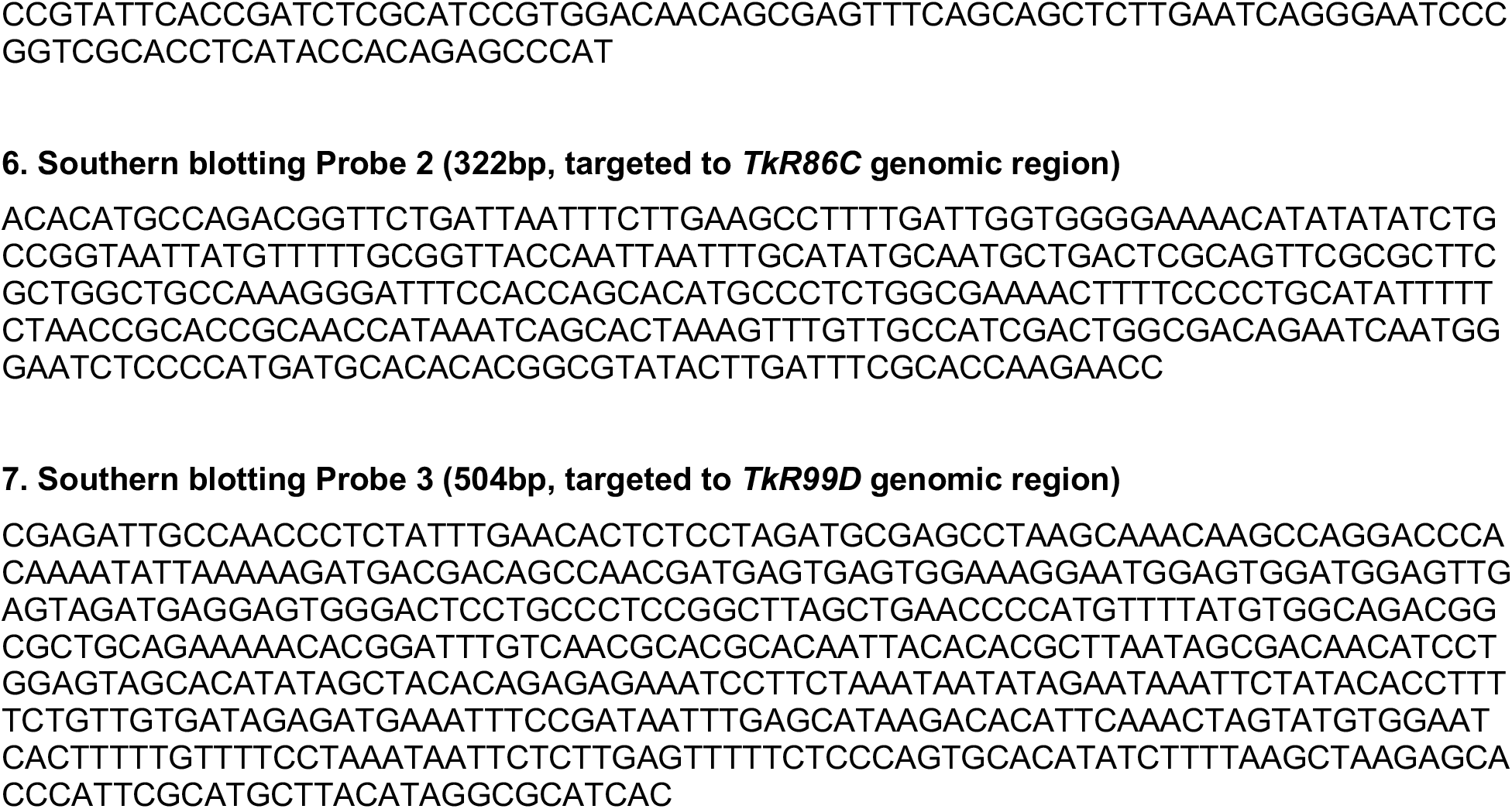
DNA sequences used in this study, related to all Figures.

